# Murine Vaginal Co-infection with Penicillinase-Producing *Neisseria gonorrhoeae* Fails to Alleviate Amoxicillin-Induced Chlamydial Persistence

**DOI:** 10.1101/2022.12.22.521649

**Authors:** Delia Onorini, Cory Ann Leonard, Theresa Pesch, Barbara Prähauser, Robert V. Schoborg, Nicole Borel

**Author notes:** denotes equal contribution, with author order decided randomly (Delia Onorini, Cory Ann Leonard).

## Abstract

*Chlamydia trachomatis* (CT) and *Neisseria gonorrhoeae* (NG) cause most bacterial sexually transmitted infections (STIs) worldwide. CT/NG co-infection is more common than expected due to chance, suggesting CT/NG interaction. However, CT/NG co-infection remains largely unstudied. Obligate intracellular CT has a characteristic biphasic developmental cycle consisting of two bacterial forms, infectious elementary bodies (EBs) and non-infectious, replicating reticulate bodies (RBs), which reside within host-derived, membrane-bound intracellular inclusions. Diverse stressors cause divergence from the normal chlamydial developmental cycle to an aberrant state called chlamydial persistence. Persistence can be induced by host-specific factors such as intracellular nutrient deprivation or cytokine exposure, and exogenous factors such as beta-lactam exposure, which disrupts RB to EB conversion. Persistent chlamydiae are atypical in appearance and, as such, are called aberrant bodies (ABs), but remain viable. The primary hallmark of persistence is reversibility of this temporary non-infectious state; upon removal of the stressor, persistent chlamydiae re-enter normal development, and production of infectious EBs resumes. The beta-lactam amoxicillin (AMX) has been shown to induce chlamydial persistence in a murine vaginal infection model, using the mouse pathogen *C. muridarum* (CM) to model human CT infection. This remains, to date, the sole experimentally tractable *in vivo* model of chlamydial persistence. Recently, we found that penicillinase-producing NG (PPNG) can alleviate AMX-induced CT and CM persistence *in vitro.* We hypothesized that PPNG vaginal co-infection would also alleviate AMX-induced CM persistence in mice. To evaluate this hypothesis, we modified the CM/AMX persistence mouse model, incorporating CM/PPNG co-infection. Contradicting our hypothesis, and recent *in vitro* findings, PPNG vaginal co-infection failed to alleviate AMX-induced CM persistence.

## INTRODUCTION

The bacterial pathogens *Chlamydia trachomatis* (CT) and *Neisseria gonorrhoeae* (NG) target the epithelium, causing sexually transmitted infections (STIs). In 2020 alone, 129 million new cases of chlamydia and 82 million new cases of gonorrhea were reported, representing most bacterial STIs reported worldwide [1], as typical for these two STIs throughout recent decades. Obligate human pathogens, CT and NG cause cervicitis in women and urethritis in men, frequently also occurring as asymptomatic and/or extragenital infections which may remain undetected and untreated. CT treatment failure is common [2], [3], and ever-increasing NG anti-microbial resistance is widespread. No vaccines are currently available for either pathogen, and the development of new CT and NG therapies and vaccines remains a significant public health concern [4], [5].

Inside the target host cells, obligate intracellular CT progresses through a characteristic bi-phasic developmental cycle [6], [7], alternating between infectious elementary bodies (EBs) and replicative reticulate bodies (RBs). Upon infection, EBs are internalized in host cell membrane-derived, *Chlamydia*-modified inclusions where they differentiate into RBs. RBs subsequently undergo several rounds of replication and differentiate again into EBs, which are then released by host cell lysis or extrusion of the intact inclusion. A variety of stressors cause divergence from the normal CT developmental cycle to an aberrant state called chlamydial persistence, or the chlamydial stress response [8]–[11]. Persistent chlamydial inclusions and bacteria are atypical in appearance but are still viable, despite stalled development which fails to progress normally to the release of infectious EBs.

Known host-specific factors that induce chlamydial persistence include intracellular nutrient deprivation or cytokine exposure [12]–[15]. Additionally, exogenous factors such as betalactam exposure, which disrupts RB to EB conversion, cause persistence and result in distinctive enlarged aberrant bodies (ABs) [16], [17]. The primary hallmark, or distinguishing characteristic, of persistence is reversibility of the induced, temporarily non-infectious, persistent state upon removal of the stressor [10], [18]. Persistent chlamydiae thus relieved of exposure to the stressor re-enter normal development and production of infectious EBs resumes. The beta-lactam amoxicillin (AMX) induces chlamydial persistence *in vitro* [8] and in a murine vaginal infection model [19], [20], the latter of which uses the mouse pathogen *Chlamydia muridarum* (CM) to model human CT infection. The murine CM/AMX model remains, to date, the sole experimentally tractable *in vivo* model of chlamydial persistence [19], [20].

Importantly, CT/NG co-infections are more common than can be explained by chance alone [21], suggesting CT/NG interaction may increase susceptibility to, or transmission of, either or both pathogens. Additionally, clinical evidence suggests that NG co-infection may reactivate potentially persistent, undetected chlamydial infection [22], [23] via unknown mechanisms. Nonetheless, CT and NG *in vitro* and *in vivo* pathogenesis studies have focused on infections with a single pathogen, with limited evaluation of CT/NG co-infection. The sole *in vivo* CM/NG co-infection model, a mouse vaginal infection model, showed higher vaginal viable NG recovery in NG co-infected mice compared to NG-only infected mice, when NG infection was initiated during acute (actively shedding) CM infection [24]. However, in our more recent murine CM/NG vaginal co-infection study, these results were not reproduced when NG infection was initiated during latent (inapparent) CM infection (manuscript under review). Additionally, in an *in vitro* tissue culture model, NG elicited anti-chlamydial effects consistent with sphingolipid depletion of the host cells [25]. Although these co-infection studies provide evidence of *Chlamydia/NG* interactions, they fail to provide evidence that NG may exert a pro-chlamydial effect, as suggested by clinical and epidemiological studies.

We aimed to evaluate the impact of NG co-infection on persistent murine genital chlamydial infection *in vivo,* utilizing the CM/AMX mouse model of chlamydial persistence, as the first proof of concept that NG can exert a measurable pro-chlamydial effect *in vivo.* We hypothesized that penicillinase-producing NG (PPNG) can alleviate AMX-induced CM persistence in the setting of *in vivo* co-infection. This is in line with our recent *in vitro* finding that PPNG co-infection can prevent and reverse AMX-induced CT and CM persistence in cultured cervical epithelial cells when direct contact of NG with CT- or CM-infected host cells is prevented (manuscript in preparation). To address our hypothesis, we modified the CM/AMX persistence mouse model [19], [20], incorporating CM/PPNG co-infection, per the previously described mouse vaginal CM/NG co-infection [24]. In the modified model, CM-infected mice were treated with AMX to induce CM persistence, as measured by cessation of vaginal viable CM shedding, followed by vaginal co-infection with PPNG or non-penicillinase-producing NG. Contradicting our hypothesis, PPNG vaginal co-infection, like nonpenicillinase-producing NG co-infection, failed to alleviate AMX-induced CM persistence in contrast to our recent CM/AMX/PPNG *in vitro* findings.

## MATERIALS AND METHODS

### Host cells and media

LLC-MK2 cells (LLC; Rhesus monkey kidney cell line; provided by IZSLER, Brescia, Italy) were cultivated as previously described [25]. LLC culture medium consisted of Dulbecco’s MEM with Ham’s F-12 (DMEM/F-12 with no tryptophan or nicotinamide; US Biological, Salem, MA, USA), supplemented with 10% heat-inactivated fetal calf serum (BioConcept, Allschwill, Switzerland), 10 mM HEPES (N-2-hydroxyethylpiperazine-N-2-ethane sulfonic acid, Sigma-Aldrich), 1.2 mg/ml Sodium Bicarbonate (Sigma-Aldrich/MilliporeSigma, St. Louis, MO, USA), 1% MEM non-essential amino acids (MEM NEAA; 100x, Gibco), 10 μg/mL L-Tryptophan (1X, Alfa Aesar, Thermo Fisher Scientific, Waltham, MA, USA), 2.02 μg/mL Nicotinamide (Sigma-Aldrich/MilliporeSigma) and 2 mM GlutaMAX-I (Gibco, Thermo Fisher Scientific), as previously described in detail [25].

### *Chlamydia muridarum* propagation and inoculum preparation

*C. muridarum* (CM) Weiss strain (obtained from Kyle Ramsey, Midwestern University) was propagated in LLC cells, crude stocks were prepared in sucrose phosphate glutamate buffer (SPG; 218 mM sucrose (Sigma-Aldrich/MilliporeSigma); 3.76 mM KH2PO4 (Sigma-Aldrich/MilliporeSigma), 7.1 mM K2HPO4 (Sigma-Aldrich/MilliporeSigma), 5 mM GlutaMAX (Gibco, Thermo Fisher Scientific)) and aliquots of stock were stored at −80°C until use [19]. Enumeration of viable EB in crude stock was determined as inclusion forming units (IFU)/ml, as previously described [25].

### *Neisseria gonorrhoeae* propagation and inoculum preparation

*Neisseria gonorrhoeae* strain FA1090 (NG; endocervical isolate, isolated in 1983 from a probable disseminated infection [26] was provided by Magnus Unemo, School of Medical Sciences, Orebro University and cultivated on commercially available chocolate agar (Thermo Scientific Chocolate agar with Vitox; Thermo Fisher Scientific) as previously described [25]. The penicillinase-producing strain (PPNG; ATCC 31426), purchased in Kwik-Stik format (Microbiologics, St. Cloud, MN, USA), was cultured on chocolate agar supplemented to 50 μg/ml final concentration of penicillin G (Sigma-Aldrich/MilliporeSigma, St. Louis, MO, USA) by adding 200 μl of 5 mg/ml penicillin G in sterile deionized water per chocolate agar plate (~20 ml agar volume), spreading and allowing to diffuse for ~30 minutes (min) prior to storage at 4°C for up to a week.

After two laboratory passages, NG stock aliquots were frozen in glycerol freezing medium (20% glycerol (Alfa Aesar, Thermo Fisher Scientific) in Tryptic Soy Broth (BD Bacto, Thermo Fisher Scientific) and filter sterilized (0.22 μm)). Before each use, NG stock aliquots were thawed, cultured, and sub-passaged once on commercial chocolate agar and PPNG stock aliquots on 50 μg/ml penicillin G chocolate agar, prior to suspension and use for vaginal inoculation (i.e., at laboratory passage 4).

To prepare inocula, NG or PPNG colonies (18-24 hours (h)) were collected with a sterile nylon swab and suspended in sterile phosphate buffered saline (PBS, pH 7.2, 1-liter tablets; Canvax Biotech, Córdoba, Spain) to a McFarland density (McF) of approximately 2.0-3.0 (DEN-1B densitometer, Grant Instruments, Cambridgeshire, UK; 0.5-4.0 McFarland Standard Set, Pro Lab Diagnostics, Richmond Hill, ON, Canada) for NG or 2.0-5.0 for PPNG. The suspended NG or PPNG was then passed through a sterile 1.2 μm filter to remove aggregates as previously described [27], and diluted with sterile PBS to a McF of ~0.4-0.6, which we determined by trial to represent colony forming units (CFU)/ml of ~10^8^ CFU/ml. Inocula, delivered as 20 μl volume per mouse, were 1.8×10^6^ CFU/ml NG and 1.5×10^6^ CFU/ml PPNG, as determined by dilution of the filtered suspensions in sterile 0.05% saponin (Sigma-Aldrich/MilliporeSigma)/ PBS cultured on Chocolate agar for 18-24 h with bacterial colonies counted on a stereo microscope (M3; Wild Heerbrugg AG, Heerbrugg, Switzerland).

### Amoxicillin (AMX) reagent and preparation

Amoxicillin (AMX; Sandoz Pharmaceuticals AG, Risch-Rotkreuz, Switzerland; 5 g amoxicillin in 11.05 g total mass, including carrier) was prepared in water-diluted condensed sweetened milk (Federation of Migros Cooperatives, Zurich, Switzerland). AMX powder was reconstituted, per manufacturer’s instruction, with autoclaved tap water to 50 mg/ml. Reconstituted AMX was stored at 4°C for less than 14 days. Diluted condensed sweetened milk was prepared by diluting the product as purchased with autoclaved tap water to 30% final concentration of milk. Diluted milk was aliquoted and stored at −20°C until use. For AMX treatment, AMX was diluted in 30% milk to a final concentration of 1.6 mg/ml as required to deliver the desired dose of 40 μg (2 mg/kg[19]; based on estimated 20 g mouse weight) in 25 μl total volume two times per day for 10 days (Days 5-14). AMX was diluted in milk the evening prior to each day of AMX treatment and stored at 4°C for less than 24 h before use.

### Co-infection and amoxicillin treatment protocol

Six-week-old female BALB/c mice were purchased from Janvier Labs (Le Genest-Saint-Isle, France). Mice were housed at five per cage, in individually ventilated cages, with 12-hour light/dark cycle, regulated temperature (21-24°C)/humidity (35-70%), with ad libitum access to standard mouse feed (Kliba Nafag; Granovit AG, Kaiseraugst, Switzerland) and water. Mice were acclimated for 12 days with tunnels included in the cages, both for enrichment and for mouse handling as refinement to reduce stress [28], [29].

Experimental design is shown in **Figure 1**; day of the experiment is noted as “Day 0” to “Day 24”. The study comprised 60 mice evaluated in two independent experiments carried out between January and April 2022 (see **Supplemental File 1**). Mice were divided into four groups: mice infected with CM but not treated with AMX (CM); mice infected with CM and treated with AMX (CM+AMX); mice infected with CM, treated with AMX and co-infected with NG (CM+AMX+NG); mice infected with CM, treated with AMX and co-infected with PPNG (CM+AMX+PPNG). On Days 0, 1 and 2, mice were vaginally inoculated for three consecutive days with 1×10^6^ IFU of CM in 10 μl SPG to facilitate inoculation in the diestrus stage of the 4-5 day reproductive cycle, which is necessary for effective chlamydial infection [30].

**Figure 1.**
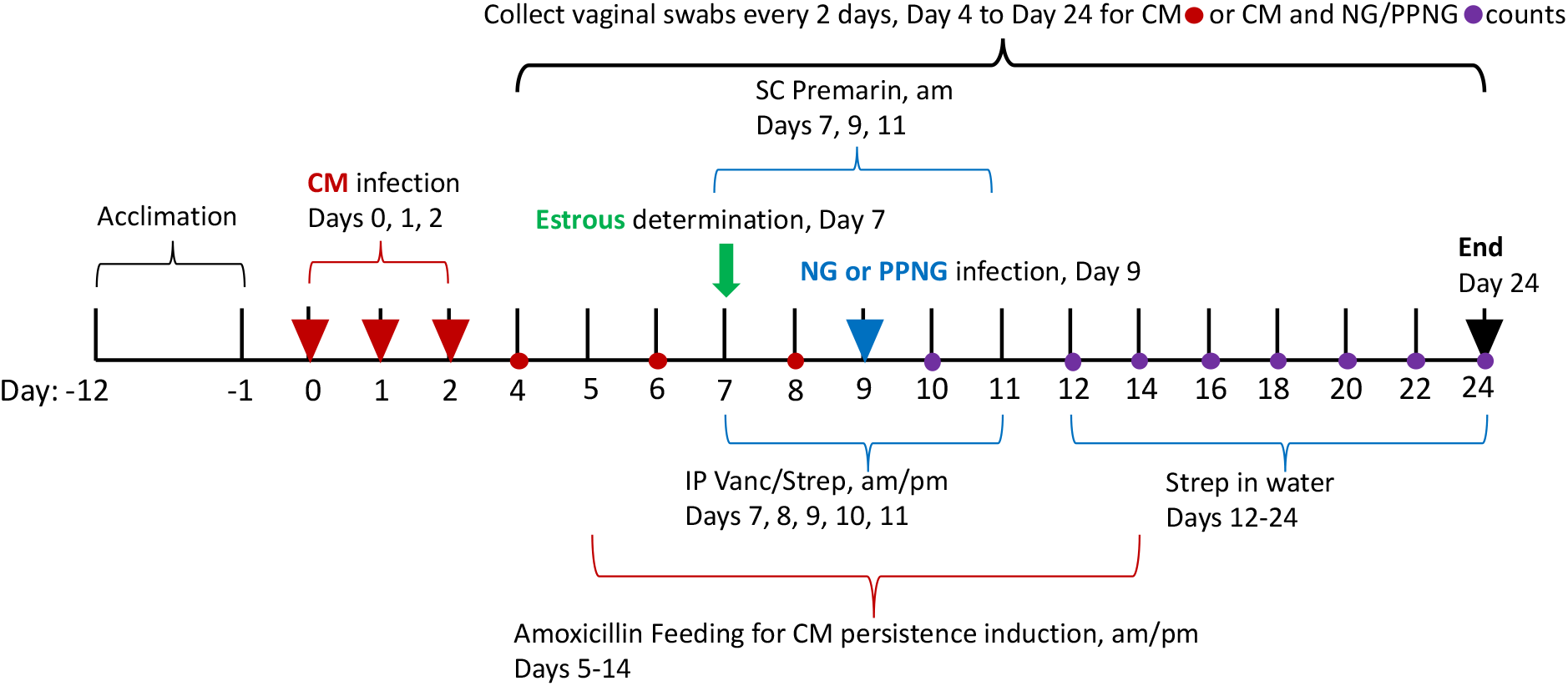
Experimental Design. Mice were vaginally inoculated with *Chlamydia muridarum* (CM) daily, for three consecutive days. Viable CM shedding was monitored by vaginal swabbing every two days from Day 4 to Day 24 of the experiment. Vaginal smears were collected on Day 7 to determine estrous cycle stage and mice in anestrus or diestrus were treated with sub-cutaneous (SC) injections of Premarin and intraperitoneal (IP) injections of antibiotics (vancomycin hydrochloride and streptomycin sulfate (Vanc/Strep) for five days, as shown. Two days after initiation of Premarin/antibiotics, mice were vaginally inoculated once with *Neisseria gonorrhoeae* (NG) or penicillinase-producing *Neisseria gonorrhoeae* (PPNG). After cessation of antibiotic injections, Strep was supplied in drinking water. Mice that were not selected for Premarin/antibiotics treatment were sacrificed on Day 7 (early sacrifice) and all other mice were sacrificed on Day 24 (late sacrifice). At sacrifice, rectal swabs were collected for viable CM enumeration and necropsy was performed to evaluate gross genital tract pathology and to collect genital and intestinal tracts tissues for pathology analyses.

On Day 5, AMX treatment was initiated and continued until Day 14 of the experiment. The twice-daily dose of 40 μg AMX was previously shown to induce CM persistence in BALB/c female mice [19]. Oral delivery of drugs via voluntary feeding in condensed sweetened milk was used to deliver the AMX in our study, as it has been previously shown to yield expected pharmacodynamics of an antipsychotic drug, while reducing measures of stress associated with oral gavage delivery [31].

On Day 7, 5 days after the last chlamydial inoculation, NG and PPNG infection was performed as previously described [24], [27]. Vaginal smear slides, prepared from all mice, were stained with Hematek Stain Pak 4481Modified Wright Stain (Siemens AG; Munich, Germany). Mice with vaginal smear cells consisting primarily of neutrophils and nucleated epithelial cells, rather than cornified epithelial cells (as determined by relative cell morphology), were considered to be in diestrus, and mice with few total cells, but visible bacteria and mucus were considered to be in anestrus[27]. Mice not in diestrus or anestrus on Day 7 were sacrificed that day. Mice in diestrus or anestrus were treated by subcutaneous injection (SC) of Premarin (0.5 mg in 200 μl, a conjugated estrogen, once per day, mornings; Sigma) on Days 7, 9 and 11 to facilitate prolonged NG colonization. Additionally, to control growth of commensal flora, caused by Premarin treatment, mice were treated by intraperitoneal injections (IP), with vancomycin hydrochloride (Vanc; 0.6 mg) and streptomycin sulfate (Strep; 2.4 mg) twice daily in 200 μl, mornings and afternoons, for five days (from Days 7-11, except Day 7, on which mice received IP injection only in the afternoon). Finally, on Days 12-24, after cessation of antibiotics injections, 5 g/l streptomycin sulfate (dissolved in autoclaved tap water) was provided ad libitum to all groups, as replacement of regular water, to continue to control commensal flora overgrowth while limiting the number of injections required.

On Day 9, approximately 4 hours after the second dose of Premarin, mice were vaginally mock inoculated with PBS (CM and CM+AMX groups) or inoculated with NG or PPNG in 20 μl of PBS. Vaginal swabs, collected every 2 days, until Day 24 (end of experiment), were evaluated first to determine vaginal chlamydial shedding of live EB (indicative of active CM infection) and chlamydial persistence (subsequent lack of detectable vaginal shedding of live EB) by titer assay. Subsequently, after NG infection, both CM shedding and recovery of viable NG from vaginal swabs were determined by titration. On Day 24, mice were sacrificed, necropsy performed, and tissue samples collected and stored for histopathology and molecular analysis. Also, upon sacrifice (on Day 7 or Day 24), rectal swabs were collected to determine CM rectal shedding.

Due to technical requirements, investigators were not blinded to any mouse procedures, sample preparations, estrous stage determination or pathology/immunohistochemistry analyses of tissues. However, CM and NG/PPNG titer analyses were done in a blinded manner.

### Sample Collection

#### Vaginal and rectal swabs, vaginal smears

Mice were swabbed (vaginally or rectally) with a PBS-soaked swab (ultra mini Flocked swab, Copan 516CS01; Copan Italia, Brescia, Italy) inserted into the vaginal canal or rectum and rotated gently. All vaginal swabs were collected from live mice, while all rectal swabs were collected from sacrificed mice. Swabs were collected into 2 ml screw-top tubes containing three sterile glass beads in 1 ml of sterile SPG. Swab tubes were then vortexed for 10 seconds and processed for recovery of viable NG (vaginal swabs only, see below) prior to snap-freezing on dry ice and subsequent storage at −80°C. Vaginal smears were similarly collected with PBS-moistened swabs, which were then smeared directly onto standard glass microscopy slides.

#### Necropsy, tissue sampling and pathology scoring

Necropsy began first with observation of gross genital tract macroscopic findings. Next, genital tract tissues (ovary, oviduct, uterus, cervix, vagina) and intestinal tract tissues (duodenum, jejunum, ileum, cecum, colon, rectum) were collected and stored in 4% buffered formaldehyde for 24 h followed by paraffin embedding (FFPE) according to routine procedure. From each FFPE block, 2 μm sections were cut and stained with hematoxylin and eosin (HE). The genital sections were subject to histologic evaluation [32] as complete longitudinal sections, by a board-certified pathologist. For assessment of the oviduct, available cross-sections were counted and the number of dilated versus total cross-sections was calculated. Additionally, presence of inflammation (yes/no) was recorded, and inflammatory infiltrates were described qualitatively (lymphocytes, macrophages and/or neutrophils) and assessed semi-quantitatively (periductal versus periovarial location for oviduct, luminal versus epithelial location for other tissues, and mild moderate or severe for degree).

#### Quantification of viable chlamydial shedding by titration and Immunofluorescence Assay (IFA) enumeration

Quantification of viable CM shedding detected in vaginal and rectal swabs was determined by titration in LLC cells. Briefly, triplicate serial dilutions were performed on thawed, vortexed swab samples used to inoculate LLC. Inocula were replaced with growth medium supplemented with 1.5 μg/ml final concentration of cycloheximide (Sigma) and antibiotics (1.3 μg/ml final concentration of amphotericin B (Gibco), 100 μg/ml final concentration of vancomycin hydrochloride (Sigma) at and 10 μg/ml final concentration of gentamycin (Gibco) and plates were incubated for 24 h at 37°C, 5% CO_2_. Cells were then fixed, and CM inclusions were immunostained with *Chlamydiaceae* mouse anti-lipopolysaccharide antibody as previously described [25]. Chlamydial inclusions were counted at 100-200X magnification, and the calculation of IFU/swab was performed (limit of detection = 5 IFU/swab, given 200 μl of total swab volume assayed across the triplicate well of the lowest sample dilution) (Nikon Eclipse TiU; Nikon Instruments Inc, Melville, NY, USA).

#### Quantification of recovery of viable NG by titration and colony enumeration

Immediately after vaginal swab collection, prior to snap freezing on dry ice, swab samples from CM+AMX+NG and CM+AMX+PPNG groups were vortexed for 5-10 seconds and added, in triplicate wells, to 0.05% saponin/PBS, in 96-well U-bottom plates and 1:3 serial dilutions (eight dilutions) were performed. Next, 5 μl of each dilution well and three 34 μl replicates of undiluted sample were plated to agar plates (Neisseria Selective Medium PLUS, Thermo Scientific) with antibiotic selection (vancomycin, colistin, amphotericin B, trimethoprim) and incubated for 24-48 h at 37°C, 5% CO_2_. NG or PPNG colonies were counted using a stereo microscope (M3; Wild Heerbrugg AG, Heerbrugg, Switzerland) and NG/PPNG recovery (CFU/swab) was calculated (limit of detection = 10 CFU/swab, given 100 μl of undiluted sample assayed) [25]. Vaginal swabs for the CM and CM+AMX groups (ie groups that were not NG-inoculated) were assayed as described, except 50 μl (undiluted) of total swab volume was assayed to confirm absence of NG/PPNG cross-contamination.

#### Tissue DNA extraction and *Chlamydiaceae* real-time PCR (qPCR)

DNA extracted from selected tissue samples (20 μm sections of FFPE blocks as previously described [33]) was eluted in 50 μl final volume. CM positivity was determined by quantitative real-time PCR based on *Chlamydiaceae* family-specific 23S rRNA gene as previously described [33], [34]. All samples were tested in duplicate with ~10^1^ or more genome copies per tissue portion detected, and the associated cycle threshold value <38 for both replicates, considered to be positive. A positive control comprising a seven-fold *C. abortus* DNA dilution series and a negative control (water instead of template DNA) were included in each run [35].

#### CM evaluation by immunohistochemistry

Complete longitudinal genital tract sections and non-longitudinal intestinal tract sections (consisting of one cross-section each from duodenum, jejunum, ileum, cecum, colon and rectum) of selected mice (n =10; CM+AMX group) were immunolabeled for CM. The primary antibody used was a *Chlamydiaceae* family-specific rabbit polyclonal antibody LPS/MOMP antibody (Cygnus Technologies, Inc., Southport, NC, USA) at a 1:1000 dilution. *Chlamydia* antigen retrieval consisted of pressure cooking (98°C) in citrate buffer (Dako/Agilent, Santa Clara, CA, USA) for 20 min. After incubation with the primary antibody for 1 h at room temperature (RT), endogenous peroxidase activity (Dako Agilent) was inhibited for 10 min at RT. Detection was performed with the Envision + System HRP Rabbit (Dako Agilent) for 30 min incubation at RT and the substrate 3-amino-9-ethylcarbazole (AEC)-peroxidase with hematoxylin counterstain. Lung tissue from a *C. pneumoniae-infected* mouse *(Chlamydia* control, kindly provided by Prof. Bernhard Kaltenboeck), was used as a positive control.

#### Mycoplasma testing

DNA extracted from cells and chlamydial stock was tested using the Mycoplasma Detection Kit for conventional PCR per manufacturer’s instructions (Venor®GeM OneStep, MB Minerva Biolabs, Germany). Cells and chlamydial stock used in this study were Mycoplasma negative.

#### Statistical analysis

Sample size analyses were performed with the statistical software R (https://www.R-project.org/). Sample size for the primary analyses of CM vaginal shedding was estimated based on simulated power analysis under design of two-way ANOVA with repeated measures model. A *final* sample size of n = 14 mice per group (considering mice dropped out due to estrous stage or undetectable PPNG colonization) was determined, to ensure the probability of 90% that the expected power is at least 0.80 to detect an effect size (signal-to-noise ratio) of 1. Experimental units were completely randomized to experimental groups in each of the independent experiments. An effect size approaching 1 was not achieved after two experiments (for Days 10-24 CM+AMX+NG vs CM+AMX+PPNG effect size, as determined by Hedges’ g test (https://www.socscistatistics.com/effectsize/default3.aspx), ranged from 0.18-0.57). Given the low effect size observed after the majority of samples (n = 9-12) had been collected in two independent experiments, an additional experiment to achieve a final sample size of n = 14 was not performed, and corresponding statistical analyses were not carried out on CM vaginal shedding, rectal shedding, or pathology outcomes.

#### 13 Animal use

All animal experiments were conducted in the Laboratory Animal Services Center (LASC) at University of Zurich (biosafety level 2) and previously approved by Cantonal Veterinarian’s Office of Zurich (License number: 018/2020). Refinements made to minimize animal stress included use of tunnels for enrichment and animal pick up (no tail pick up), alternating IP injection site left/right for daily IP injections, limiting overall number of IP injections by providing antibiotics in water as possible, and utilizing voluntary feeding of AMX in sweetened condensed milk to eliminate standard oral gavage delivery.

#### 14 Data Availability

The original contributions presented are included within the article and the Supplemental Material. Further inquiries can be directed to the corresponding author.

## RESULTS

### Study Design

We evaluated the impact of PPNG co-infection on AMX-induced persistent murine genital chlamydial infection, hypothesizing that PPNG in the setting of *in vivo* co-infection could alleviate AMX-induced CM persistence. To address our hypothesis, we modified the existing CM/AMX persistence mouse model [19], [20], incorporating CM/PPNG co-infection similar to that previously described for mouse vaginal CM/NG co-infection [24], as detailed in (**Figure 1**).

First, we vaginally inoculated all mice with CM once daily for three days. We used the same inoculum dose of 10^6^ CM IFU as is our recent latent CM/NG murine vaginal co-infection study (manuscript under review), which resulted in detectable CM infection of nearly all mice in both studies. Next, two days after the final CM inoculation, we initiated treatment of CM-infected mice with AMX, to induce CM persistence, and maintained AMX treatment for 10 days; mice not treated with AMX served as non-persistent controls (ie CM group; acutely infected). Then, two days after initiation of AMX treatment, mice in all groups were evaluated for estrous stage, and those not in anestrus or diestrus were sacrificed (“early sacrifice”), while those in anestrus/diestrus were immediately treated Premarin (estrogen) and antibiotics, via an injection regimen, to facilitate prolonged NG infection in mice subsequently infected with NG or PPNG [24], [27], [36]. Finally, two days after initiation of the Premarin/antibiotics regimen, AMX-treated mice were vaginally co-infected with NG or PPNG or were similarly mock-inoculated with NG diluent alone. The NG co-infected group served as a control for NG co-infection absent of penicillinase production. Thus, all experimental groups in the study received equivalent CM inoculation or corresponding mock inoculation, identical Premarin/antibiotic treatment, and equivalent NG or PPNG inoculation or corresponding mock inoculation.

Mice were monitored for the presence of vaginal viable CM, NG and PPNG by collection of vaginal swabs every two days. All mice not sacrificed on Day 7 were sacrificed on Day 24 (“late sacrifice”) at the end of the experiment. Rectal swabs and tissues (genital and intestinal) were collected from all sacrificed mice (early or late). We monitored vaginal viable CM shedding to confirm the rapid cessation, or near cessation, of CM vaginal shedding characteristic of induction of chlamydial persistence [19], [20], as well as to determine any effect of PPNG to alleviate this persistence. We additionally monitored recovery of viable NG and PPNG from vaginal swabs for both *N. gonorrhoeae* strains and rectal shedding of CM.

The study consisted of two independent experiments (10 or 50 mice each, 60 total mice, **Supplemental File 1**). Mice comprised four experimental groups: CM, acute chlamydial infection; CM+AMX, persistent chlamydial infection; CM+AMX+NG, persistent chlamydial infection subject to co-infection with a non-penicillinase-producing strain of *N. gonorrhoeae;* and CM+AMX+PPNG, persistent chlamydial infection subject to co-infection with a penicillinase-producing strain of *N. gonorrhoeae*. Mice not in anestrus or diestrus, and thus subject to early sacrifice, as well as mice with no detectable CM or PPNG in any vaginal swab (**Supplemental File 1**) were excluded from enumeration analyses of viable vaginal bacteria (**Figure 2**), but not from pathology analyses (**Tables 4-7**). Mice with no detectable NG in any swab were not excluded from analyses because NG is AMX susceptible, and robust, prolonged infection in the presence of AMX is not expected. Experimental group sizes after the described exclusions were n = 9 mice, for CM and CM+AMX+PPNG groups, and n = 12 mice, for CM+AMX and CM+AMX+NG groups.

**Figure 2.**
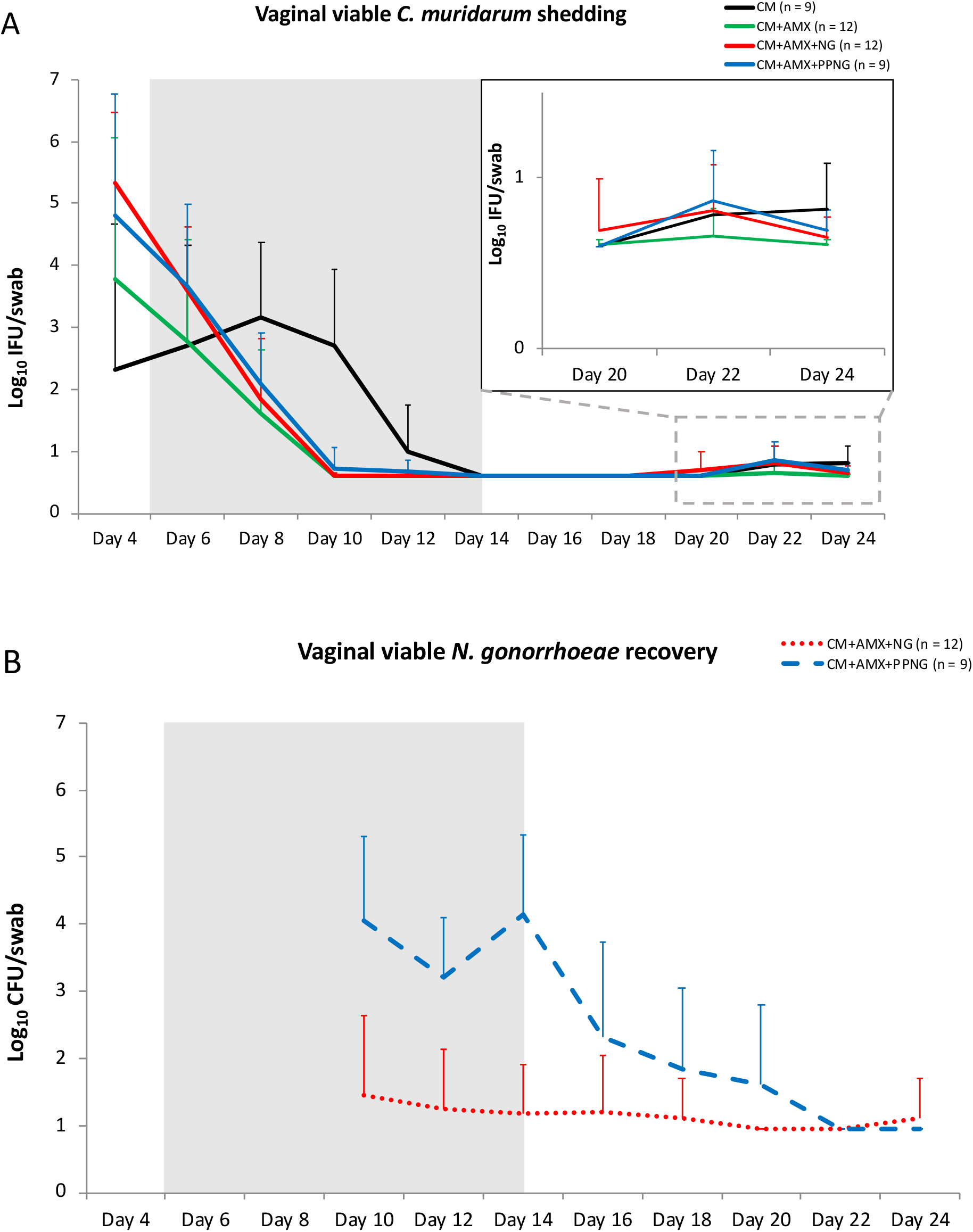
Vaginal viable *Chlamydia muridarum* (CM) shedding and recovery of vaginal viable *Neisseria gonorrhoeae* (NG) and penicillinase-producing NG (PPNG). Viable CM shedding (**A**) was determined by infecting LLC cells with vaginal swab medium, while viable NG recovery (**B**) was determined by culturing vaginal swab medium on agar plates selective for *N. gonorrhoeae.* Results are expressed as mean viable bacteria per swab: log_10_ inclusion forming units (IFU)/swab for CM and log_10_ colony forming units (CFU)/swab for NG and PPNG (+ standard deviation). CM vaginal inoculation was done on Days 0, 1 and 2 of the experiment, NG and PPNG vaginal inoculation was done on Day 9, and AMX treatment was done from Day 5-14 (grey shading). Early sacrifice mice and mice with no detectable CM or PPNG, but not NG, in any swab were excluded from analysis. For swabs with no detectable shedding, the CM IFU/swab was set at 4 (ie log_10_ = 0.60) because the CM titer limit of detection is 5, while the IFU/swab was set at 9 (ie log_10_ = 0.95) because the NG and PPNG titer limit of detection is 10.

In the first independent experiment of the study (Experiment 1, n = 10 mice, CM+AMX group; see **Supplemental File 1**), we evaluated CM genome copy number *(Chlamydia* qPCR, CM qPCR) and CM immunohistochemical (IHC) positivity for all formalin-fixed genital and intestinal tissues collected from AMX-treated early sacrifice mice (n = 4) and late sacrifice mice (n = 6). Of the four early sacrifice mice, three were positive for CM in both genital and intestinal tissue sections (log_10_ CM genomes = 1.60-2.65), and one was positive in only the intestine (log_10_ CM genomes = 2.70); of the six late sacrifice mice, two were positive in the intestine only (log_10_ CM genome copies of 0.93-1.65). However, by IHC analysis, no CM inclusions could be detected in the genital or intestinal epithelia of these mice, regardless of CM qPCR positivity. This contrasts findings from a recent study carried out on similarly CM-infected, but non-AMX-treated, mice sacrificed at Day 23, wherein >50% of CM-infected mice had occasional intestinal inclusions (manuscript under review). We speculated that, in the current study, CM tissue inclusions were undetected due to AMX treatment and/or due to the relatively early timepoint for the early sacrifice mice. Tissue-associated CM qPCR or IHC analyses were not carried out for the subsequent experiment of the study.

### Vaginal co-infection with penicillinase-producing *N. gonorrhoeae* (PPNG) does not alleviate amoxicillin-induced *C. muridarum* (CM) persistence

We hypothesized that reduced vaginal viable CM shedding, the primary outcome evaluated in this study, would be alleviated by PPNG co-infection, resulting in increased CM shedding in the CM+AMX+PPNG group compared to that of the CM+AMX and CM+AMX+NG groups. Vaginal viable CM shedding was assessed by quantitative titer assay (culture in LLC host cells) and analyzed and presented as mean “CM shedding” in log_10_ CM IFU/swab (SD, standard deviation) (**Figure 2 A**). Vaginal viable NG or PPNG recovery was assessed by quantitative titer assay (in selective agar) and analyzed and presented as mean “NG recovery” in log_10_ NG CFU/swab (SD) (**Figure 2 B**). Additionally, proportion/percent of mice with detectable CM shedding and NG or PPNG recovery are shown in **Table 1** and **Table 2**, respectively. Finally, raw CM shedding data and NG/PPNG recovery data are provided in **Supplemental File 2** and **Supplemental File 3**, respectively.

**Table 1.** Vaginal swab *Chlamydia muridarum* titer^1^.

| Table 1. Vaginal swab *Chlamydia muridarum* titer<sup>1</sup> |  |  |  |  |  |  |  |  |  |  |  |
| --- | --- | --- | --- | --- | --- | --- | --- | --- | --- | --- | --- |
| Group | Day 4 | Day 6 | Day 8 | Day 10 | Day 12 | Day 14 | Day 16 | Day 18 | Day 20 | Day 22 | Day 24 |
| CM | 5/9<br>(56%) | 7/9<br>(78%) | 9/9<br>(100%) | 8/9<br>(89%) | 4/9<br>(44%) | 0/9 | 0/9 | 0/9 | 0/9 | 5/9<br>(56%) | 5/9<br>(56%) |
| CM +AMX | 11/12<br>(92%) | 10/12<br>(83%) | 9/12<br>(75%) | 0/12 | 0/12 | 1/12<br>(8%) | 0/12 | 0/12 | 1/12<br>(8%) | 2/12<br>(17%) | 1/12<br>(8%) |
| CM +AMX +NG | 12/12<br>(100%) | 12/12<br>(100%) | 9/12<br>(75%) | 2/12<br>(17%) | 0/12 | 0/12 | 0/12 | 0/12 | 1/12<br>(8%) | 5/12<br>(42%) | 3/12<br>(25%) |
| CM +AMX +PPNG | 9/9<br>(100%) | 8/9<br>(89%) | 8/9<br>(89%) | 1/9<br>(11%) | 1/9<br>(11%) | 0/9 | 0/9 | 0/9 | 0/9 | 5/9<br>(56%) | 5/9<br>(56%) |
| 1 Positive/total (percent) of mice with vaginal viable CM shedding<br>Early sacrifice mice and mice with no detectable CM, NG or PPNG in any swab were excluded from analysis |  |  |  |  |  |  |  |  |  |  |  |

**Table 2.** Vaginal swab *Neisseria gonorrhoeae* titer^1^.

| Table 2. Vaginal swab *Neisseria gonorrhoeae* titer<sup>1</sup> |  |  |  |  |  |  |  |  |  |  |  |
| --- | --- | --- | --- | --- | --- | --- | --- | --- | --- | --- | --- |
| Group | Day 4 | Day 6 | Day 8 | Day 10 | Day 12 | Day 14 | Day 16 | Day 18 | Day 20 | Day 22 | Day 24 |
| CM +AMX +NG | NA | NA | NA | 2/12<br>(17%) | 2/12<br>(17%) | 1/12<br>(8%) | 1/12<br>(8%) | 1/12<br>(8%) | 0/12 | 0/12 | 1/12<br>(8%) |
| CM +AMX +PPNG | NA | NA | NA | 9/9<br>(100%) | 9/9<br>(100%) | 9/9<br>(100%) | 5/9<br>(56%) | 4/9<br>(44%) | 3/9<br>(33%) | 0/9 | 0/9 |
| 1 Positive/total (percent) of mice with vaginal viable NG or PPNG recovered<br>NA = not applicable; Day 4, 6 and 9 vaginal swabs were collected prior to NG or PPNG inoculation<br>Early sacrifice mice and mice with no detectable CM, NG or PPNG in any swab were excluded from analysis |  |  |  |  |  |  |  |  |  |  |  |

From Day 4-24 of the study, as expected, control acutely CM-infected mice (not exposed to AMX) exhibited peak shedding consistent with that described for acute vaginal shedding, namely peak vaginal shedding 6-9 days after inoculation, followed by decline to undetectable, or near undetectable, levels of viable CM shedding, as occurs during natural progression from acute infection to CM latency [30], [37]. Specifically, at peak CM shedding on Day 8, mean log_10_ (SD) IFU/swab for the CM group was 3.16 (1.21) (**Figure 2 A**). Decline to latency (no/low detection) occurred by Day 14, with no shedding detected from Days 14-20, and a low level of shedding again detected on Days 22 and 24, with mean shedding of 0.81 (0.27) and 0.65 (0.12), respectively. The proportion of mice with detectable CM shedding in the CM control group also peaked at Day 8 (100%), with a range of 44% to 100% positivity from Days 4-12; the proportion of mice with detectable CM shedding on Days 22 and 24 was 56% (**Table 1**). Day 4 vaginal viable CM shedding across all early sacrifice mice (n = 11, 4.20 (2.19)) versus all late sacrifice mice (n = 49, 4.01 (1.48)) was similar prior to initiation of AMX treatment.

In contrast, as expected, AMX treatment resulted in a sharp drop in detectable vaginal viable CM shedding. AMX treated groups, CM+AMX, CM+AMX+NG, and CM+AMX+PPNG, had Day 4 shedding values of 3.77 (2.30), 5.33 (1.14), and 4.81 (1.95), respectively (**Figure 2 A**); 92-100% of mice in AMX treated groups were positive for CM shedding on Day 4 (**Table 1**). These AMX treated groups all had very limited proportions of mice with CM shedding detected from Days 10-20, ranging from 8-17% positivity across all three groups; on Days 22 and 24, mean log_10_ (SD) IFU/swab did not exceed 0.86 (0.31) for any of the AMX treated groups (**Figure 2 A**). Proportions of mice with resumed detectable CM shedding on Days 22 and 24 were 8-17% for the CM+AMX group, 25-42% for the CM+AMX+NG group and 56% for the CM+AMX+PPNG group (**Table 1**); Day 22 or 24 CM shedding values for individual mice ranged from the limit of detection, 0.70, to 1.40 across all four groups of the study. While these comparisons are speculative due to small group sizes, the observed proportion of mice positive for CM shedding is potentially larger for PPNG co-infected versus NG co-infected mice.

We evaluated vaginal viable NG and PPNG recovery, to determine if, and to what degree, mouse/vaginal swab CM shedding positivity on Days 10-24 was concurrent with NG and/or PPNG recovery. Recovery of vaginal viable NG, which does not produce penicillinase, was detected in 8/96 (8%) swabs analyzed for the CM+AMX+NG group (**Supplemental File 3**) and mean NG log_10_ CFU/swab ranged from 1.12-1.45, when detected, on Days 10-24 (**Figure 2 B**). Of the 96 swabs evaluated for the CM+AMX+NG group, 12 (13%) had detectable CM shedding; however, only 1/12 (8%) of these CM-positive swabs (from Day 10) were also NG recovery positive. And, while the NG recovery value for this CM- and NG-positive mouse was the highest detected in the study (4.57), the CM shedding value for this mouse was the lowest detectable CM shedding value, at the limit of CM detection (0.70). Thus, as expected, detectable vaginal viable non-penicillinase-producing NG recovery does not appear to impact CM shedding during AMX-induced persistence.

Recovery of viable PPNG, which produces penicillinase was detected in 39/72 (54%) swabs analyzed for the CM+AMX+PPNG group (**Supplemental File 3**) and mean PPNG recovery values ranged from 1.59-5.49, when detected, on Days 10-20 (no PPNG was recovered on Days 22 and 24) (**Figure 2 B**). Of the 72 swabs evaluated for the CM+AMX+PPNG group, 12 (17%) had detectable CM shedding; however, only 2/12 (17%) of the CM-positive swabs, from a single mouse on Days 10 and 12, were also PPNG recovery positive. The PPNG recovery values for this mouse on Day 10 and Day 12 swabs was 3.68 and 3.98, respectively. However, the remaining 8 mice in the group also had PPNG recovery on Days 10 and 12, ranging in value from 1.69-5.49, but had no detectable CM shedding. Additionally, the 10 CM+AMX+PPNG swabs positive for CM shedding on Days 22 and 24 had no associated detectable PPNG recovery. Thus, we find no clear evidence, in contrast to our expectation, that detectable vaginal viable PPNG impacts CM shedding during AMX-induced persistence.

### Vaginal infection with *C. muridarum* (CM) is associated with rectal carriage of viable CM in the presence or absence of AMX treatment

Vaginal CM inoculation has been consistently associated with both genital tract, and subsequent intestinal tract, viable CM carriage [38]–[41]. Rectal viable CM shedding was assessed by quantitative titer assay (culture in LLC host cells) and analyzed and presented as percent of mice with detectable CM rectal shedding, and as mean group CM shedding in log_10_ CM IFU/swab (SD) (**Table 3**). Of 11 rectal swabs collected at early sacrifice, the sole CM group early sacrifice mouse was not rectal swab CM positive; while 60% of CM-infected, AMX-treated early sacrifice mice were rectal swab CM positive (mean shedding = 1.54 (1.00)). Of 22 rectal swabs collected at late sacrifice, CM and CM+AMX groups had similar percent of mice with detectable rectal CM shedding, 67% and 62%, respectively; CM and CM+AMX groups also had similar mean rectal shedding, 1.49 (1.12) and 1.46 (1.09), respectively. Thus, we find no evidence that AMX treatment alone modulates late sacrifice rectal viable CM positivity. Interestingly, of 27 rectal swabs collected at late sacrifice, CM+AMX+NG and CM+AMX+PPNG groups had lower rectal viable CM shedding values than the CM or CM+AMX groups. CM+AMX+NG and CM+AMX+PPNG groups had a similar percent of mice with detectable rectal shedding, 25% and 27%, respectively, and had similar mean log_10_ CM IFU/ rectal swab (SD), 1.02 (0.86) and 0.97 (0.76), respectively. This leads us to speculate that *N. gonorrhoeae* co-infection may negatively affect rectal carriage of viable CM.

**Table 3.** Rectal swab *Chlamydia muridarum* titer^1^.

| Table 3. Rectal swab *Chlamydia muridarum* titer<sup>1</sup> |  |  |
| --- | --- | --- |
| Group | Early sacrifice | Late sacrifice |
| CM | 0/1<br>0.60 | 6/9 (67%)<br>1.49 (1.22) |
| CM+AMX | 3/7 (71%)<br>0.99 (0.56) | 8/13 (62%)<br>1.46 (1.09) |
| CM+AMX+NG | 3/3 (100%)<br>2.82 (0.22) | 3/12 (25%)<br>1.02 (0.86) |
| CM+AMX+PPNG | None | 4/15 (27%)<br>0.97 (0.76) |
| ALL CM+AMX | 6/10 (60%) | NA |
| 1 Positive/total (percent) mice with rectal viable CM shedding; mean log<sub>10</sub> CM IFU/swab (standard deviation)<br>NA = not applicable; early sacrifice, but not late sacrifice, AMX-treated groups are equivalent<br>None = CM+AMX+PPNG group had no early sacrifice mice<br>For swabs with no detectable CM, IFU/swab was set at 4 (ie log<sub>10</sub> = 0.60); CM titer limit of detection is 5 |  |  |

### Pathology Analyses of the Genital Tract

For all early and late sacrifice mice, across all groups, we observed no remarkable gross genital tract pathology. To facilitate pathological assessment of the full genital tract (longitudinal section), two sections were made from each fixed genital tract block for each mouse (n = 60; 11 early sacrifice and 49 late sacrifice). The ovary, present in at least one section for all mice, was similar in normal appearance across all experimental groups and was thus not further evaluated. The oviduct, uterus, cervix and vagina were evaluated for dilation and the presence of inflammatory cells.

At least one oviduct cross section was present for all mice. Oviduct cross sections/mouse ranged from 1 to 12 (mean = 5.22, 2.44 SD). Oviduct dilation was observed in only two mice (**Table 4; Supplemental File 4**), one early sacrifice (CM+AMX) and one late sacrifice (CM+AMX+PPNG) mouse. Oviduct inflammation, was not observed in either mouse having oviduct dilation, but was present in 16 mice (**Table 5; Supplemental File 4**). In early sacrifice mice, oviduct inflammation positivity ranged from 57-100% across all groups, while in late sacrifice mice positivity ranged from 7-33% across all groups. In line with the expected association of CM infection with oviduct pathology [42], [43], both early and late sacrifice mice without AMX treatment appeared to have potentially higher proportions of mice with oviduct inflammation. For early sacrifice mice, oviduct inflammation was observed in 100% of non-AMX-treated mice, and 57-67% of AMX-treated mice; while for late sacrifice mice, oviduct inflammation was observed in 33% of non-AMX-treated mice, and 7-25% of AMX-treated mice. However, these comparisons must be regarded as speculative due to small group sizes.

**Table 4.**
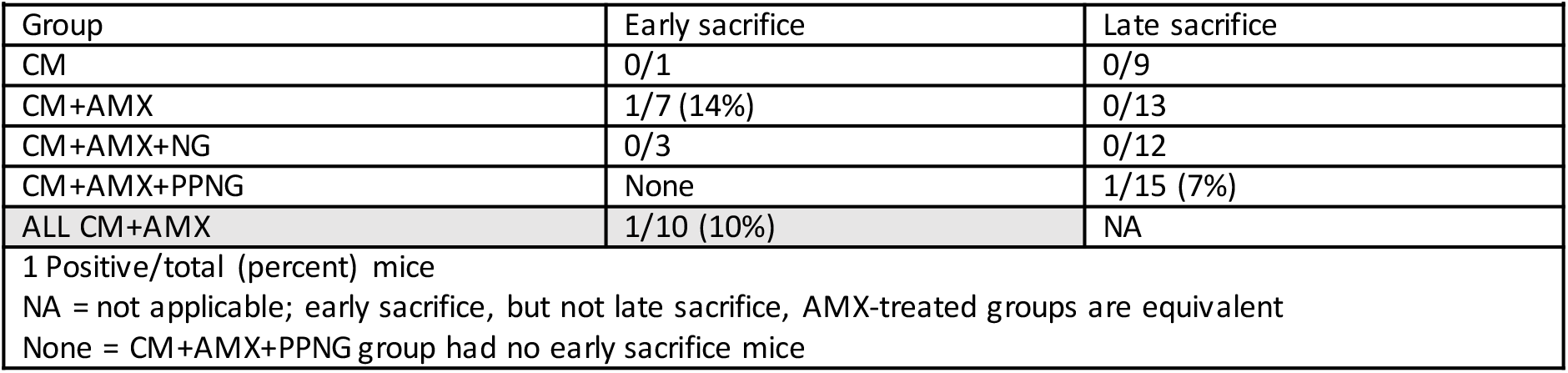
Oviduct Dilation^1^.

| **Table 4. Oviduct Dilation<sup>1</sup>** |  |  |
| --- | --- | --- |
| Group | Early sacrifice | Late sacrifice |
| CM | 0/1 | 0/9 |
| CM+AMX | 1/7 (14%) | 0/13 |
| CM+AMX+NG | 0/3 | 0/12 |
| CM+AMX+PPNG | None | 1/15 (7%) |
| ALL CM+AMX | 1/10 (10%) | NA |
| 1 Positive/total (percent) mice<br>NA = not applicable; early sacrifice, but not late sacrifice, AMX-treated groups are equivalent<br>None = CM+AMX+PPNG group had no early sacrifice mice |  |  |

**Table 5.** Oviduct Inflammation^1^.

| **Table 5. Oviduct Inflammation<sup>1</sup>** |  |  |
| --- | --- | --- |
| Group | Early sacrifice | Late sacrifice |
| CM | 1/1 (100%) | 3/9 (33%) |
| CM+AMX | 4/7 (57%) | 2/13 (15%) |
| CM+AMX+NG | 2/3 (67%) | 3/12 (25%) |
| CM+AMX+PPNG | None | 1/15 (7%) |
| ALL CM+AMX | 6/10 (60%) | NA |
| 1 Positive/total (percent) mice<br>NA = not applicable; early sacrifice, but not late sacrifice, AMX-treated groups are equivalent<br>None = CM+AMX+PPNG group had no early sacrifice mice |  |  |

Of the mice with oviduct inflammation, 63% had inflammatory cells only in periductal areas, and most of these mice (80%) were in the late sacrifice group. The remaining mice (38%) with oviduct inflammation had both periductal and periovarial inflammatory cell localization; of these mice, most (83%) were in the early sacrifice group. Inflammatory cell types present in the oviduct included macrophages and neutrophils. Early sacrifice mice had exclusively multiple types of inflammatory cells type present (all three cell types in five mice, and lymphocytes and macrophages in one mouse), while most late sacrifice mice (70%) had only lymphocytes present. Focal distribution of oviduct inflammatory cells was observed only in late sacrifice mice, and was primarily associated with lymphocyte-only inflammatory cells.

Tissue was available for pathological analyses of the uterus in all mice (**Supplemental File 4**), but we do not speculate pathology or infection-associated instance for these observations. Additionally, we expect Premarin treatment to facilitate an estrus-like state, but we also expect that CM and/or NG/PPNG infection may potentially confound tissue-based analysis of estrus stage, due to potential neutrophil infiltration. In line with this, genital tissues of nearly all mice at early and late sacrifice were consistent with estrus or proestrus/estrus stages (not shown).

The cervix, present for 58 mice, was lined by columnar epithelium transitioning to stratified epithelium (**Supplemental File 4; Table 6**). Presence of cervical inflammatory cells (exclusively neutrophils) was relatively common. For early sacrifice mice, the sole CM group mouse had cervical inflammation, while 20% of the AMX-treated mice (all in the CM+AMX group) had cervical inflammation. Cervical inflammation rates were generally higher for late sacrifice mice; 25% of CM mice, 38% of CM+AMX mice, 58% of CM+AMX+NG mice and 21% of CM+AMX+PPNG mice had cervical inflammation. Cervical inflammation was detected in 20 mice. In all cases, intraepithelial infiltration was present, more commonly as intraepithelial infiltration alone (n = 13) than as combined intraepithelial infiltration with luminal accumulation (n = 7). Accumulation/infiltration ranged from mild to severe and typically, except in one case, did not differ between luminal and intraepithelial sites when present at both.

**Table 6.** Cervical Inflammation^1^.

| **Table 6. Cervical Inflammation<sup>1</sup>** |  |  |
| --- | --- | --- |
| Group | Early sacrifice | Late sacrifice |
| CM | 1/1 (100%) | 2/8 (25%)* |
| CM+AMX | 2/7 (29%) | 5/13 (38%) |
| CM+AMX+NG | 0/3 | 7/12 (58%) |
| CM+AMX+PPNG | None | 3/14 (21%)* |
| ALL CM+AMX | 2/10 (20%) | NA |
| 1 Positive/total (percent) mice<br>*One mouse in the group is missing the cervical section<br>NA = not applicable; early sacrifice, but not late sacrifice, AMX-treated groups are equivalent<br>None = CM+AMX+PPNG group had no early sacrifice mice |  |  |

The vagina, present in sections of 55 mice, was primarily (n = 42, 76%) lined with keratinized squamous epithelial cells, or, less frequently (n = 13, 24%), with stratified epithelial cells (**Supplemental File 4; Table 7**). Presence of vaginal inflammatory cells (exclusively neutrophils) was common. For early sacrifice mice with vaginal sections present, CM+AMX and CM+AMX+NG groups, 71% and 67% of mice had vaginal inflammation, respectively. These rates were higher than those observed for late sacrifice mice, which ranged from 25-58%, amongst which CM+AMX+NG and CM+AMX+PPNG groups both had the same vaginal inflammation rate (58%). Vaginal inflammation, detected in 29 mice, was most common as intraepithelial infiltration alone (n = 22, 76%), less common as combined intraepithelial infiltration with luminal accumulation (n = 5, 17%), and relatively uncommon (n = 2, 7%) as luminal accumulation only. Accumulation/infiltration ranged from mild to severe across all mice, an unlike cervical inflammation, generally differed between luminal and intraepithelial sites when present at both. Of the 29 mice positive for vaginal inflammation, 10 (34%) were positive for cervical inflammation and 6 (21%) for oviduct inflammation (**Supplemental File 4**).

**Table 7.** Vaginal inflammation^1^.

| **Table 7. Vaginal inflammation<sup>1</sup>** |  |  |
| --- | --- | --- |
| Group | Early sacrifice | Late sacrifice |
| CM | * | **2/8 (25%) |
| CM+AMX | 5/7 (71%) | 6/13 (46%) |
| CM+AMX+NG | 2/3 (67%) | 7/12 (58%) |
| CM+AMX+PPNG | None | ***7/12 (58%) |
| ALL CM+AMX | 7/10 (70%) | NA |
| 1 Positive/total (percent) mice<br>*The single mouse in the group is missing the vaginal section<br>**One mouse in the group is missing the vaginal section<br>***Three mice in the group are missing vaginal sections<br>NA = not applicable; early sacrifice, but not late sacrifice, AMX-treated groups are equivalent<br>None = CM+AMX+PPNG group had no early sacrifice mice |  |  |

## DISCUSSION

Potential mechanisms by which CT/NG interaction may increase host susceptibility, host transmission of infection, or both, include increased bacterial load of one or both pathogens and/or increased duration of viable pathogen shedding. NG co-infection may reactivate undetected female genital chlamydial shedding [23], [44], [45], possibly via one of these mechanisms. However, the two existing CM/NG murine vaginal co-infection studies to date, have thus far not provided evidence of such a “pro-chlamydial” effect of NG co-infection on existing CM murine vaginal infection. The first published murine vaginal CM/NG co-infection study showed that NG co-infection of acutely CM-infected mice, with active vaginal viable CM shedding of approximately 2.0-3.0 log_10_ IFU/swab, had no effect on CM shedding; while recovery of vaginal viable NG (approximately 2.0-3.0 log_10_ CFU/swab) was significantly increased, by about 0.5-1.0 log_10_ CFU/swab, though duration of NG shedding was not influenced [24]. More recently, we found that similar vaginal NG-coinfection of CM-infected mice, when delayed to the naturally later-occurring latent stage of CM infection, as characterized by undetectable CM shedding, failed to elicit such an effect on viable NG recovery, as well as having no effect on viable CM shedding magnitude or duration (manuscript under review). Thus, we aimed herein to evaluate the potential of a penicillinase-producing strain of NG (PPNG) to alleviate AMX-induced (beta lactam-induced) CM persistence in a mouse vaginal infection model, considering that this effect, as recently observed in an *in vitro* setting in our laboratory (manuscript in preparation) may provide the first, and thus far elusive, proof of concept that NG may play a measurable pro-chlamydial role *in vivo.*

To enable exploration of our hypothesis, we modified the only experimentally tractable chlamydial persistence mouse model [19], [20], by incorporating CM/PPNG co-infection, per the existing mouse vaginal CM/NG co-infection [24]. In our modified model, AMX-induction of CM persistence, was observable as expected, as rapid cessation of vaginal viable CM shedding as previously described [19], [20]. Furthermore, despite the presumed potential for different estrous stage in early versus late sacrifice mice at the initiation of CM inoculations, given the 4-5 day cycle [46] and the early sacrifice of *only* non-anestrus/diestrus mice, mean log_10_ (SD) IFU/swab was similar in early and late sacrifice mice. Thus, we conclude that three consecutive days of CM inoculation has relatively equivalent CM infection outcome regardless of initial estrous stage, and represents an effective approach for CM vaginal infection mouse studies. These observations indicate that the model as employed herein is appropriate and sufficient, as far as is currently possible, to evaluate our hypothesis.

Nonetheless, contradicting our hypothesis (and in contrast to our recent CM/AMX/PPNG *in vitro* findings; manuscript in preparation), PPNG vaginal co-infection had no notable effect on AMX-induced CM persistence – neither increasing magnitude of vaginal viable CM shedding, nor markedly increasing the proportion of mice with vaginal viable CM shedding. Additionally, PPNG recovery was not typically associated with the instances of post-PPNG-infection CM shedding within the CM+AMX+PPNG group. Our findings suggest that 1) actively replicating PPNG does not express sufficient penicillinase to cleave the relevant AMX molecules, 2) and/or that PPNG growth is not sufficient to provide and alternative AMX target sufficient to limit the effect of AMX on CM in the *in vivo* setting, or 3) that PPNG-produced penicillinase is degraded or inactivated in our *in vivo* setting. Given that further observations in a continued study designed to have n = 14 group sizes (relative to the n = 9-12 group sizes already achieved), would not feasibly be sufficient to result in the planned effect size of 1.0, with the planned/allotted total number of mice, we chose to discontinue the study to eliminate further animal use. We thus conclude, qualitatively, that we observed no effect of PPNG co-infection on persistent CM vaginal infection in mice.

Vaginal, intragastric and rectal CM inoculation have been shown to result in sustained intestinal infection [38]–[41]. Interestingly, such rectal infection is less susceptible to cure with azithromycin treatment (the primary recommended CT therapy, [47]) than concomitant vaginal infection in mice, as measured by detection of surviving viable CM in cecal, but not cervical, tissues despite similar azithromycin levels in cervical and cecal tissues [48]. Clinical studies have similarly reported rectal CT azithromycin treatment failures in men and women [49]–[51]. In line with these previous mouse study and clinical findings, our current findings suggest that the effect of AMX on CM may also be limited in the setting of the intestinal tract, relative to the genital tract, at least in mice.

Though statistical analysis was not performed, the NG and PPNG co-infected groups had a rectal viable CM positivity rate value approximately half that of the non-co-infected groups, and a mean log_10_ CM IFU/ rectal swab value approximately two-thirds that of the non-co-infected groups. This suggests that NG or PPNG co-infection of CM-infected, AMX-treated mice may be associated with lower magnitude and/or proportion of rectal CM carriage – a potential effect not explained by NG strain specificity or penicillinase production. Notably, this is consistent with our recently published findings that NG co-infection of epithelial host cells *in vitro* (in human genital and intestinal cell lines) has an anti-chlamydial effect on CT and CM infection, reducing inclusion size, numbers and infectivity [25]. Thus, anti-chlamydial effects of NG may occur both *in vitro* and *in vivo,* prompting us to consider how such effects might be reconciled with the high rates of clinical CT/NG co-infection and the expectation of synergistic, not competitive, CT/NG bacterial interaction. We may speculate that reduced intensity of chlamydial infection may be associated with longer duration of chlamydial infection in humans. In line with this possibility, it has been shown that a lower vaginal inoculation dose of CM (single CM inoculation after progesterone treatment) was associated with increased numbers of detectable viable chlamydiae in the murine oviduct [52].

Gross genital tract observations in our progesterone-free co-infection model, as observed here and in our recent similar study (manuscript under review), showed no hydrosalpinx, as typically observed for progesterone-enhanced CM murine vaginal infection [53], [54]. Progesterone-free CM vaginal infection by others also showed no hydrosalpinx [55], so less gross genital tract pathology may be characteristic of hormone-free CM infection. We also assessed pathology 22 days after the final CM inoculation, and genital pathology measures are often reported later, up 80 days post-infection [53]. Additionally, relatively high CM inoculum as we used (10^5^-10^7^ IFU) has been shown to result in *less* ascending infection and hydrosalpinx [52]. Finally, stress has been shown to increase CM-associated pathology [56], [57], so our use of tunnels may have reduced stress, compared to standard tail-handling experiments [28], [29], which may in turn be speculated to reduce overall CM-associated pathology.

Oviduct pathology is typical of murine CM infection. In line with this, we found that oviduct inflammation may be less prevalent in AMX-treated groups. Oviduct inflammation was also potentially more prevalent at early sacrifice versus late sacrifice timepoints. However, because sacrifice early after CM infection was restricted to mice not in diestrus/anestrus, it is not possible to differentiate between the potential impact of early timepoint and non-diestrus/anestrus stage on oviduct inflammation. Neither NG nor PPNG co-infection clearly increased oviduct dilation or inflammation. Oviduct inflammation rates may be mildly increased in the CM+AMX+NG group compared to the CM+AMX and CM+AMX+PPNG late sacrifice groups. However, this contrasts previous findings for NG co-infection during CM latency (using the same strains of CM and NG used herein), which did not increase oviduct dilation or inflammation (manuscript under review). Finally, in the current study, oviduct inflammation rates at late sacrifice (Day 24, 7-33%) are relatively similar to those in the previous latency study at early or late sacrifice (Day 23, 25% and Day 36, 4-27%, respectively). However, mice in the current study have notably less oviduct dilation (<4% of all mice, across all groups) than our previous latency study, which ranged from about 25-40% across all CM-infected groups. This might be due to the shorter study duration or AMX-treatment in the current study; however, both mice with oviduct dilation in the current study were in AMX-treated groups, which does not support the likelihood of this possibility.

In contrast to oviduct inflammation, cervical and vaginal inflammation did not appear to be limited, or less pronounced, in mice treated with AMX. NG co-infection was associated with the highest observed prevalence of both cervical and vaginal inflammation across all late sacrifice groups, but any role of NG infection in cervical pathology is speculative due to small sample size. Similarly, in our previous latent CM/NG mouse vaginal co-infection study (manuscript under review) we observed no specific association of CM and/or NG infection with cervical or vaginal inflammation, and no increase in inflammation for co-infected mice. We expect the presence of inflammatory cells, especially in the cervix and vagina, may be impacted by the requirements of our model system, such as frequent vaginal manipulations (inoculation, swabbing) or potential effects of antibiotics on vaginal normal flora.

In summary, we 1) show that PPNG co-infection does not appear to reactivate CM vaginal shedding which has been supressed by AMX-induced CM persistence, 2) postulate that the ability of AMX to supress viable CM shedding may be limited in the intestinal tract, compared to the genital tract, 3) propose that relatively minimal oviduct pathology may be associated with progesterone-free CM infection, and 4) find no clear evidence that NG/PPNG coinfection causes, or is associated with, remarkable genital pathology.

A major limitation of our study, though currently no alternative has been reported, is the requirement for estrogen treatment to facilitate successful, sustained NG infection – given that an induced state of estrus may be unfavorable for CM genital infection, which is better supported by the diestrus/anestrus stage [30]. Similar estradiol treatment in the original CM/NG co-infection model did not prevent vaginal viable CM shedding during acute vaginal CM infection [24]. However, we cannot rule out a negative effect of estrogen treatment on CM infection as a potential contributing factor in the failure of PPNG co-infection to alleviate CM persistence. Pilot studies to determine if vaginally-delivered intact/active exogenous penicillinase can measurably alleviate AMX-induced CM persistence may be warranted prior to further utilization of the CM/AMX/PPNG co-infection model. Despite limitations of current CM/NG co-infection mouse models, however, continued *in vivo* evaluation of *Chlamydia/NG* interaction, is important to inform ongoing efforts toward novel and improved prevention and treatment strategies for chlamydia and gonorrhea.

## ACKNOWLEDGEMENTS

We thank the molecular laboratory team of the Institute of Veterinary Pathology, Zurich (IVPZ) and Dagmar Schäfer and the University of Zurich Laboratory Animal Services Center (LASC) team for technical assistance with tissue samples. We also thank Professor Ann Jerse for valuable discussion regarding the murine CM/NG co-infection model. Finally, we would like to thank Giacomo Marziali for his contributions to sample collection, processing and evaluation.

