## Supplemental File 1 for "Murine Vaginal Co-infection with Penicillinase-Producing *Neisseria gonorrhoeae* Fails to Alleviate Amoxicillin-Induced Chlamydial Persistence"

### SUPPLEMENTAL FILE 1. Study Design

| Expt | Group | Cage | Mouse | Early Sac | Status |
|---|---|---|---|---|---|
| 1 | CM+AMX | 1 | 1 | Yes | ND |
| 1 | CM+AMX | 1 | 2 | No | OK |
| 1 | CM+AMX | 1 | 3 | No | OK |
| 1 | CM+AMX | 1 | 4 | Yes | ND |
| 1 | CM+AMX | 1 | 5 | No | OK |
| 1 | CM+AMX | 2 | 6 | Yes | ND |
| 1 | CM+AMX | 2 | 7 | No | OK |
| 1 | CM+AMX | 2 | 8 | Yes | ND |
| 1 | CM+AMX | 2 | 9 | No | OK |
| 1 | CM+AMX | 2 | 10 | No | NCM |

…..

| Expt | Group | Cage | Mouse | Early Sac | Status |
|---|---|---|---|---|---|
| 2 | CM | 1 | 1 | Yes | ND |
| 2 | CM | 1 | 2 | No | OK |
| 2 | CM | 1 | 3 | No | OK |
| 2 | CM | 1 | 4 | No | OK |
| 2 | CM | 1 | 5 | No | OK |
| 2 | CM | 2 | 6 | No | OK |
| 2 | CM | 2 | 7 | No | OK |
| 2 | CM | 2 | 8 | No | OK |
| 2 | CM | 2 | 9 | No | OK |
| 2 | CM | 2 | 10 | No | OK |
| 2 | CM+AMX | 3 | 11 | No | OK |
| 2 | CM+AMX | 3 | 12 | No | OK |
| 2 | CM+AMX | 3 | 13 | No | OK |
| 2 | CM+AMX | 3 | 14 | No | OK |
| 2 | CM+AMX | 3 | 15 | Yes | ND |
| 2 | CM+AMX | 4 | 16 | No | OK |
| 2 | CM+AMX | 4 | 17 | No | OK |
| 2 | CM+AMX | 4 | 18 | No | OK |
| 2 | CM+AMX | 4 | 19 | Yes | ND |
| 2 | CM+AMX | 4 | 20 | Yes | ND |
| 2 | CM+AMX+NG | 1 | 1 | No | OK |
| 2 | CM+AMX+NG | 1 | 2 | Yes | ND |
| 2 | CM+AMX+NG | 1 | 3 | No | NNG |
| 2 | CM+AMX+NG | 1 | 4 | No | NNG |
| 2 | CM+AMX+NG | 1 | 5 | No | NNG |
| 2 | CM+AMX+NG | 2 | 6 | No | NNG |
| 2 | CM+AMX+NG | 2 | 7 | No | NNG |
| 2 | CM+AMX+NG | 2 | 8 | Yes | ND |
| 2 | CM+AMX+NG | 2 | 9 | No | NNG |
| 2 | CM+AMX+NG | 2 | 10 | No | OK |
| 2 | CM+AMX+NG | 3 | 11 | No | NNG |
| 2 | CM+AMX+NG | 3 | 12 | No | NNG |
| 2 | CM+AMX+NG | 3 | 13 | Yes | ND |
| 2 | CM+AMX+NG | 3 | 14 | No | NNG |
| 2 | CM+AMX+NG | 3 | 15 | No | OK |
| 2 | CM+AMX+PPNG | 1 | 1 | No | OK |
| 2 | CM+AMX+PPNG | 1 | 2 | No | OK |
| 2 | CM+AMX+PPNG | 1 | 3 | No | OK |
| 2 | CM+AMX+PPNG | 1 | 4 | No | OK |
| 2 | CM+AMX+PPNG | 1 | 5 | No | OK |
| 2 | CM+AMX+PPNG | 2 | 6 | No | NNG |
| 2 | CM+AMX+PPNG | 2 | 7 | No | NNG |
| 2 | CM+AMX+PPNG | 2 | 8 | No | NNG |
| 2 | CM+AMX+PPNG | 2 | 9 | No | OK |
| 2 | CM+AMX+PPNG | 2 | 10 | No | OK |
| 2 | CM+AMX+PPNG | 3 | 11 | No | OK |
| 2 | CM+AMX+PPNG | 3 | 12 | No | NNG |
| 2 | CM+AMX+PPNG | 3 | 13 | No | OK |
| 2 | CM+AMX+PPNG | 3 | 14 | No | NNG |
| 2 | CM+AMX+PPNG | 3 | 15 | No | NNG |

**Key:** CM = *Chlamydia muridarum*, AMX = Amoxicillin, NG = *Neisseria gonorrhoeae* (strain FA1090, does not produce Penicillinase), PPNG = Penicillinase-producing *Neisseria gonorrhoeae*. Experimental Groups: CM (CM-infected, not AMX-treated, not NG- or PPNG-infected); CM+AMX (CM-infected and AMX-treated, not NG- or PPNG-infected); CM+AMX+NG (CM-infected and AMX-treated, NG co-infected); CM+AMX+PPNG (CM-infected and AMX-treated, PPNG co-infected). Early Sac = sacrificed on Day 7 (Yes or No). Status: ND = not in Diestrus/Anestrus on Day 7; OK = in Diestrus/Anestrus and successfully CM and/or NG infected (included in vaginal bacterial shedding analyses); NCM = in Diestrus/Anestrus on Day 7, but CM infection not detected (excluded from vaginal bacterial shedding analyses); NNG = in Diestrus/Anestrus on Day 7, but NG or PPNG infection not detected (NNG status CM+AMX+PPNG mice were excluded from vaginal bacterial shedding analysis).
