## Supplemental File 2 for "Murine Vaginal Co-infection with Penicillinase-Producing *Neisseria gonorrhoeae* Fails to Alleviate Amoxicillin-Induced Chlamydial Persistence"

### SUPPLEMENTAL FILE 2. Raw vaginal viable CM shedding data

| Expt | Group | Cage | Mouse | Early Sac | Status | Day 4 | Day 6 | Day 8 | Day 10 | Day 12 | Day 14 | Day 16 | Day 18 | Day 20 | Day 22 | Day 24 |
|---|---|---|---|---|---|---|---|---|---|---|---|---|---|---|---|---|
| 2 | CM | 1 | 1 | Yes | ND | 4.92 | 4.60 | | | | | | | | | |
| 2 | CM | 1 | 2 | No | OK | 1.18 | 3.53 | 3.29 | 3.23 | 0.60 | 0.60 | 0.60 | 0.60 | 0.60 | 0.70 | 1.18 |
| 2 | CM | 1 | 3 | No | OK | 0.60 | 0.60 | 1.48 | 1.85 | 0.60 | 0.60 | 0.60 | 0.60 | 0.60 | 1.40 | 0.60 |
| 2 | CM | 1 | 4 | No | OK | 2.97 | 0.60 | 2.53 | 0.60 | 0.60 | 0.60 | 0.60 | 0.60 | 0.60 | 0.60 | 0.60 |
| 2 | CM | 1 | 5 | No | OK | 7.03 | 5.53 | 4.76 | 2.69 | 0.60 | 0.60 | 0.60 | 0.60 | 0.60 | 0.60 | 0.60 |
| 2 | CM | 2 | 6 | No | OK | 5.28 | 1.74 | 4.85 | 3.69 | 2.44 | 0.60 | 0.60 | 0.60 | 0.60 | 1.18 | 1.18 |
| 2 | CM | 2 | 7 | No | OK | 0.60 | 3.40 | 1.54 | 3.41 | 0.70 | 0.60 | 0.60 | 0.60 | 0.60 | 0.70 | 1.18 |
| 2 | CM | 2 | 8 | No | OK | 1.95 | 3.01 | 3.75 | 3.67 | 0.70 | 0.60 | 0.60 | 0.60 | 0.60 | 0.60 | 0.60 |
| 2 | CM | 2 | 9 | No | OK | 0.60 | 3.92 | 3.13 | 4.06 | 2.20 | 0.60 | 0.60 | 0.60 | 0.60 | 0.60 | 0.70 |
| 2 | CM | 2 | 10 | No | OK | 0.60 | 2.08 | 3.13 | 1.18 | 0.60 | 0.60 | 0.60 | 0.60 | 0.60 | 0.70 | 0.70 |

| Expt | Group | Cage | Mouse | Early Sac | Status | Day 4 | Day 6 | Day 8 | Day 10 | Day 12 | Day 14 | Day 16 | Day 18 | Day 20 | Day 22 | Day 24 |
|---|---|---|---|---|---|---|---|---|---|---|---|---|---|---|---|---|
| 1 | CM+AMX | 1 | 1 | Yes | ND | 4.07 | 3.18 | | | | | | | | | |
| 1 | CM+AMX | 1 | 2 | No | OK | 3.92 | 2.92 | 0.60 | 0.60 | 0.60 | 0.60 | 0.60 | 0.60 | 0.60 | 0.60 | 0.60 |
| 1 | CM+AMX | 1 | 3 | No | OK | 1.85 | 0.60 | 0.60 | 0.60 | 0.60 | 0.60 | 0.60 | 0.60 | 0.60 | 0.60 | 0.60 |
| 1 | CM+AMX | 1 | 4 | Yes | ND | 0.60 | 1.54 | | | | | | | | | |
| 1 | CM+AMX | 1 | 5 | No | OK | 7.08 | 5.53 | 1.54 | 0.60 | 0.60 | 0.60 | 0.60 | 0.60 | 0.60 | 0.60 | 0.60 |
| 1 | CM+AMX | 2 | 6 | Yes | ND | 5.05 | 3.27 | | | | | | | | | |
| 1 | CM+AMX | 2 | 7 | No | OK | 0.70 | 2.16 | 0.70 | 0.60 | 0.60 | 0.60 | 0.60 | 0.60 | 0.70 | 0.60 | 0.60 |
| 1 | CM+AMX | 2 | 8 | Yes | ND | 4.43 | 2.70 | | | | | | | | | |
| 1 | CM+AMX | 2 | 9 | No | OK | 4.27 | 1.54 | 1.18 | 0.60 | 0.60 | 0.60 | 0.60 | 0.60 | 0.60 | 0.60 | 0.60 |
| 1 | CM+AMX | 2 | 10 | No | NCM | 0.60 | 0.60 | 0.60 | 0.60 | 0.60 | 0.60 | 0.60 | 0.60 | 0.60 | 0.60 | 0.60 |
| 2 | CM+AMX | 3 | 11 | No | OK | 0.60 | 3.56 | 0.60 | 0.60 | 0.60 | 0.60 | 0.60 | 0.60 | 0.60 | 0.70 | 0.60 |
| 2 | CM+AMX | 3 | 12 | No | OK | 4.67 | 2.66 | 2.22 | 0.60 | 0.60 | 0.60 | 0.60 | 0.60 | 0.60 | 0.60 | 0.60 |
| 2 | CM+AMX | 3 | 13 | No | OK | 3.58 | 3.73 | 3.35 | 0.60 | 0.60 | 0.60 | 0.60 | 0.60 | 0.60 | 0.60 | 0.60 |
| 2 | CM+AMX | 3 | 14 | No | OK | 1.81 | 1.00 | 3.57 | 0.60 | 0.60 | 0.60 | 0.60 | 0.60 | 0.60 | 0.60 | 0.60 |
| 2 | CM+AMX | 3 | 15 | Yes | ND | 5.65 | 3.41 | | | | | | | | | |
| 2 | CM+AMX | 4 | 16 | No | OK | 6.75 | 4.48 | 1.81 | 0.60 | 0.60 | 0.70 | 0.60 | 0.60 | 0.60 | 0.60 | 0.60 |
| 2 | CM+AMX | 4 | 17 | No | OK | 3.13 | 0.60 | 1.00 | 0.60 | 0.60 | 0.60 | 0.60 | 0.60 | 0.60 | 1.18 | 0.70 |
| 2 | CM+AMX | 4 | 18 | No | OK | 6.88 | 4.58 | 2.04 | 0.60 | 0.60 | 0.60 | 0.60 | 0.60 | 0.60 | 0.60 | 0.60 |
| 2 | CM+AMX | 4 | 19 | Yes | ND | 4.04 | 2.48 | | | | | | | | | |
| 2 | CM+AMX | 4 | 20 | Yes | ND | 3.96 | 3.10 | | | | | | | | | |

| Expt | Group | Cage | Mouse | Early Sac | Status | Day 4 | Day 6 | Day 8 | Day 10 | Day 12 | Day 14 | Day 16 | Day 18 | Day 20 | Day 22 | Day 24 |
|---|---|---|---|---|---|---|---|---|---|---|---|---|---|---|---|---|
| 2 | CM+AMX+NG | 1 | 1 | No | OK | 4.32 | 3.05 | 1.48 | 0.60 | 0.60 | 0.60 | 0.60 | 0.60 | 0.60 | 1.18 | 0.60 |
| 2 | CM+AMX+NG | 1 | 2 | Yes | ND | 2.34 | 1.00 | | | | | | | | | |
| 2 | CM+AMX+NG | 1 | 3 | No | NNG | 6.13 | 3.14 | 0.60 | 0.60 | 0.60 | 0.60 | 0.60 | 0.60 | 0.60 | 1.18 | 1.00 |
| 2 | CM+AMX+NG | 1 | 4 | No | NNG | 4.84 | 3.13 | 2.45 | 0.60 | 0.60 | 0.60 | 0.60 | 0.60 | 0.60 | 0.60 | 0.70 |
| 2 | CM+AMX+NG | 1 | 5 | No | NNG | 3.95 | 1.74 | 3.08 | 0.60 | 0.60 | 0.60 | 0.60 | 0.60 | 0.60 | 0.60 | 0.60 |
| 2 | CM+AMX+NG | 2 | 6 | No | NNG | 5.34 | 3.71 | 2.33 | 0.60 | 0.60 | 0.60 | 0.60 | 0.60 | 0.60 | 0.60 | 0.60 |
| 2 | CM+AMX+NG | 2 | 7 | No | NNG | 6.57 | 4.44 | 2.48 | 0.60 | 0.60 | 0.60 | 0.60 | 0.60 | 0.60 | 0.60 | 0.60 |
| 2 | CM+AMX+NG | 2 | 8 | Yes | ND | 3.49 | 1.88 | | | | | | | | | |
| 2 | CM+AMX+NG | 2 | 9 | No | NNG | 3.92 | 2.15 | 0.60 | 0.60 | 0.60 | 0.60 | 0.60 | 0.60 | 1.65 | 0.60 | 0.60 |
| 2 | CM+AMX+NG | 2 | 10 | No | OK | 3.90 | 3.41 | 3.39 | 0.70 | 0.60 | 0.60 | 0.60 | 0.60 | 0.60 | 1.18 | 0.60 |
| 2 | CM+AMX+NG | 3 | 11 | No | NNG | 7.04 | 5.24 | 2.26 | 0.60 | 0.60 | 0.60 | 0.60 | 0.60 | 0.60 | 0.60 | 0.60 |

| | | | | | | | | | | | | | | | | |
|---|---|---|---|---|---|---|---|---|---|---|---|---|---|---|---|---|
| 2 | CM+AMX+NG | 3 | 12 | No | NNG | 6.09 | 4.33 | 1.00 | 0.60 | 0.60 | 0.60 | 0.60 | 0.60 | 0.60 | 1.00 | 0.60 |
| 2 | CM+AMX+NG | 3 | 13 | Yes | ND | 5.61 | 2.50 | | | | | | | | | |
| 2 | CM+AMX+NG | 3 | 14 | No | NNG | 5.34 | 4.22 | 0.60 | 0.70 | 0.60 | 0.60 | 0.60 | 0.60 | 0.60 | 0.60 | 0.60 |
| 2 | CM+AMX+NG | 3 | 15 | No | OK | 6.50 | 4.63 | 1.93 | 0.60 | 0.60 | 0.60 | 0.60 | 0.60 | 0.60 | 1.00 | 0.70 |

| Expt | Group | Cage | Mouse | Early Sac | Status | Day 4 | Day 6 | Day 8 | Day 10 | Day 12 | Day 14 | Day 16 | Day 18 | Day 20 | Day 22 | Day 24 |
|---|---|---|---|---|---|---|---|---|---|---|---|---|---|---|---|---|
| 2 | CM+AMX+PPNG | 1 | 1 | No | OK | 2.80 | 3.55 | 2.54 | 1.65 | 1.18 | 0.60 | 0.60 | 0.60 | 0.60 | 1.00 | 0.60 |
| 2 | CM+AMX+PPNG | 1 | 2 | No | OK | 6.03 | 3.97 | 2.66 | 0.60 | 0.60 | 0.60 | 0.60 | 0.60 | 0.60 | 1.30 | 0.70 |
| 2 | CM+AMX+PPNG | 1 | 3 | No | OK | 6.60 | 4.62 | 2.13 | 0.60 | 0.60 | 0.60 | 0.60 | 0.60 | 0.60 | 0.60 | 0.60 |
| 2 | CM+AMX+PPNG | 1 | 4 | No | OK | 5.02 | 3.79 | 3.27 | 0.60 | 0.60 | 0.60 | 0.60 | 0.60 | 0.60 | 0.60 | 0.60 |
| 2 | CM+AMX+PPNG | 1 | 5 | No | OK | 6.56 | 4.52 | 2.20 | 0.60 | 0.60 | 0.60 | 0.60 | 0.60 | 0.60 | 1.18 | 0.70 |
| 2 | CM+AMX+PPNG | 2 | 6 | No | NNG | 7.10 | 5.17 | 1.90 | 0.60 | 0.60 | 0.60 | 0.60 | 0.60 | 0.60 | 0.60 | 0.60 |
| 2 | CM+AMX+PPNG | 2 | 7 | No | NNG | 4.65 | 2.55 | 3.27 | 0.70 | 0.60 | 0.60 | 0.60 | 0.60 | 0.60 | 0.60 | 0.60 |
| 2 | CM+AMX+PPNG | 2 | 8 | No | NNG | 4.27 | 2.64 | 0.60 | 0.60 | 0.60 | 0.60 | 0.60 | 0.60 | 0.60 | 1.30 | 1.00 |
| 2 | CM+AMX+PPNG | 2 | 9 | No | OK | 5.54 | 4.21 | 2.38 | 0.60 | 0.60 | 0.60 | 0.60 | 0.60 | 0.60 | 1.18 | 1.00 |
| 2 | CM+AMX+PPNG | 2 | 10 | No | OK | 2.75 | 2.64 | 1.00 | 0.60 | 0.60 | 0.60 | 0.60 | 0.60 | 0.60 | 0.60 | 0.60 |
| 2 | CM+AMX+PPNG | 3 | 11 | No | OK | 6.47 | 5.02 | 2.06 | 0.60 | 0.60 | 0.60 | 0.60 | 0.60 | 0.60 | 0.60 | 0.70 |
| 2 | CM+AMX+PPNG | 3 | 12 | No | NNG | 6.55 | 4.64 | 2.84 | 0.60 | 0.60 | 0.60 | 0.60 | 0.60 | 0.60 | 1.00 | 0.60 |
| 2 | CM+AMX+PPNG | 3 | 13 | No | OK | 1.48 | 0.60 | 0.60 | 0.60 | 0.60 | 0.60 | 0.60 | 0.60 | 0.60 | 0.70 | 0.70 |
| 2 | CM+AMX+PPNG | 3 | 14 | No | NNG | 3.52 | 4.32 | 1.00 | 0.60 | 0.60 | 0.60 | 0.60 | 0.60 | 0.60 | 0.60 | 0.60 |
| 2 | CM+AMX+PPNG | 3 | 15 | No | NNG | 5.67 | 4.12 | 2.79 | 0.60 | 0.60 | 0.60 | 0.60 | 0.60 | 0.60 | 0.60 | 0.70 |

**Key:** CM = *Chlamydia muridarum*, AMX = Amoxicillin, NG = *Neisseria gonorrhoeae* (strain FA1090, does not produce Penicillinase), PPNG = Penicillinase-producing *Neisseria gonorrhoeae*. Experimental Groups: CM (CM-infected, not AMX-treated, not NG- or PPNG-infected); CM+AMX (CM-infected and AMX-treated, not NG- or PPNG-infected); CM+AMX+NG (CM-infected and AMX-treated, NG co-infected); CM+AMX+PPNG (CM-infected and AMX-treated, PPNG co-infected). Early Sac = sacrificed on Day 7 (Yes or No). Status: ND = not in Diestrus/Anestrus on Day 7; OK = in Diestrus/Anestrus and successfully CM and/or NG infected (included in vaginal bacterial shedding analyses); NCM = in Diestrus/Anestrus on Day 7, but CM infection not detected (excluded from vaginal bacterial shedding analyses); NNG = in Diestrus/Anestrus on Day 7, but NG or PPNG infection not detected (NNG status CM+AMX+PPNG mice were excluded from vaginal bacterial shedding analysis). Day 4-24 CM shedding values are shown as the log$_{10}$ of the calculated IFU/swab; limit of detection = 5, so undetectable shedding was designated as 4 IFU/swab (i.e. <5) with corresponding log$_{10}$ value of 0.60 (indicated by red text). Shaded values are those excluded from vaginal viable CM shedding analysis shown in Figure 2 due to ND, NCM or NNG status. Where no value is entered for a given Day, the mouse was sacrificed early, and thus no longer in the study on that Day.
