## Supplemental File 3 for "Murine Vaginal Co-infection with Penicillinase-Producing *Neisseria gonorrhoeae* Fails to Alleviate Amoxicillin-Induced Chlamydial Persistence"

### SUPPLEMENTAL FILE 3. Raw vaginal viable NG and PPNG recovery data

| Expt | Group | Cage | Mouse | Early Sac | Status | Day 10 | Day 12 | Day 14 | Day 16 | Day 18 | Day 20 | Day 22 | Day 24 |
|---|---|---|---|---|---|---|---|---|---|---|---|---|---|
| 2 | CM | 1 | 1 | Yes | ND | | | | | | | | |
| 2 | CM | 1 | 2 | No | OK | none | none | none | none | none | none | none | none |
| 2 | CM | 1 | 3 | No | OK | none | none | none | none | none | none | none | none |
| 2 | CM | 1 | 4 | No | OK | none | none | none | none | none | none | none | none |
| 2 | CM | 1 | 5 | No | OK | none | none | none | none | none | none | none | none |
| 2 | CM | 2 | 6 | No | OK | none | none | none | none | none | none | none | none |
| 2 | CM | 2 | 7 | No | OK | none | none | none | none | none | none | none | none |
| 2 | CM | 2 | 8 | No | OK | none | none | none | none | none | none | none | none |
| 2 | CM | 2 | 9 | No | OK | none | none | none | none | none | none | none | none |
| 2 | CM | 2 | 10 | No | OK | none | none | none | none | none | none | none | none |

| Expt | Group | Cage | Mouse | Early Sac | Status | Day 10 | Day 12 | Day 14 | Day 16 | Day 18 | Day 20 | Day 22 | Day 24 |
|---|---|---|---|---|---|---|---|---|---|---|---|---|---|
| 1 | CM+AMX | 1 | 1 | Yes | ND | | | | | | | | |
| 1 | CM+AMX | 1 | 2 | No | OK | none | none | none | none | none | none | none | none |
| 1 | CM+AMX | 1 | 3 | No | OK | none | none | none | none | none | none | none | none |
| 1 | CM+AMX | 1 | 4 | Yes | ND | | | | | | | | |
| 1 | CM+AMX | 1 | 5 | No | OK | none | none | none | none | none | none | none | none |
| 1 | CM+AMX | 2 | 6 | Yes | ND | | | | | | | | |
| 1 | CM+AMX | 2 | 7 | No | OK | none | none | none | none | none | none | none | none |
| 1 | CM+AMX | 2 | 8 | Yes | ND | | | | | | | | |
| 1 | CM+AMX | 2 | 9 | No | OK | none | none | none | none | none | none | none | none |
| 1 | CM+AMX | 2 | 10 | No | NCM | none | none | none | none | none | none | none | none |
| 2 | CM+AMX | 3 | 11 | No | OK | none | none | none | none | none | none | none | none |
| 2 | CM+AMX | 3 | 12 | No | OK | none | none | none | none | none | none | none | none |
| 2 | CM+AMX | 3 | 13 | No | OK | none | none | none | none | none | none | none | none |
| 2 | CM+AMX | 3 | 14 | No | OK | none | none | none | none | none | none | none | none |
| 2 | CM+AMX | 3 | 15 | Yes | ND | | | | | | | | |
| 2 | CM+AMX | 4 | 16 | No | OK | none | none | none | none | none | none | none | none |
| 2 | CM+AMX | 4 | 17 | No | OK | none | none | none | none | none | none | none | none |
| 2 | CM+AMX | 4 | 18 | No | OK | none | none | none | none | none | none | none | none |
| 2 | CM+AMX | 4 | 19 | Yes | ND | | | | | | | | |
| 2 | CM+AMX | 4 | 20 | Yes | ND | | | | | | | | |

| Expt | Group | Cage | Mouse | Early Sac | Status | Day 10 | Day 12 | Day 14 | Day 16 | Day 18 | Day 20 | Day 22 | Day 24 |
|---|---|---|---|---|---|---|---|---|---|---|---|---|---|
| 2 | CM+AMX+NG | 1 | 1 | No | OK | 3.26 | 0.95 | 0.95 | 3.91 | 2.95 | 0.95 | 0.95 | 2.96 |
| 2 | CM+AMX+NG | 1 | 2 | Yes | ND | | | | | | | | |
| 2 | CM+AMX+NG | 1 | 3 | No | NNG | 0.95 | 0.95 | 0.95 | 0.95 | 0.95 | 0.95 | 0.95 | 0.95 |
| 2 | CM+AMX+NG | 1 | 4 | No | NNG | 0.95 | 0.95 | 0.95 | 0.95 | 0.95 | 0.95 | 0.95 | 0.95 |
| 2 | CM+AMX+NG | 1 | 5 | No | NNG | 0.95 | 0.95 | 0.95 | 0.95 | 0.95 | 0.95 | 0.95 | 0.95 |
| 2 | CM+AMX+NG | 2 | 6 | No | NNG | 0.95 | 0.95 | 0.95 | 0.95 | 0.95 | 0.95 | 0.95 | 0.95 |
| 2 | CM+AMX+NG | 2 | 7 | No | NNG | 0.95 | 0.95 | 0.95 | 0.95 | 0.95 | 0.95 | 0.95 | 0.95 |
| 2 | CM+AMX+NG | 2 | 8 | Yes | ND | | | | | | | | |
| 2 | CM+AMX+NG | 2 | 9 | No | NNG | 0.95 | 0.95 | 0.95 | 0.95 | 0.95 | 0.95 | 0.95 | 0.95 |
| 2 | CM+AMX+NG | 2 | 10 | No | OK | 4.57 | 4.06 | 3.52 | 0.95 | 0.95 | 0.95 | 0.95 | 0.95 |
| 2 | CM+AMX+NG | 3 | 11 | No | NNG | 0.95 | 0.95 | 0.95 | 0.95 | 0.95 | 0.95 | 0.95 | 0.95 |

| | | | | | | | | | | | | | |
|---|---|---|---|---|---|---|---|---|---|---|---|---|---|
| 2 | CM+AMX+NG | 3 | 12 | No | NNG | 0.95 | 0.95 | 0.95 | 0.95 | 0.95 | 0.95 | 0.95 | 0.95 |
| 2 | CM+AMX+NG | 3 | 13 | Yes | ND | | | | | | | | |
| 2 | CM+AMX+NG | 3 | 14 | No | NNG | 0.95 | 0.95 | 0.95 | 0.95 | 0.95 | 0.95 | 0.95 | 0.95 |
| 2 | CM+AMX+NG | 3 | 15 | No | OK | 0.95 | 1.29 | 0.95 | 0.95 | 0.95 | 0.95 | 0.95 | 0.95 |

| Expt | Group | Cage | Mouse | Early Sac | Status | Day 10 | Day 12 | Day 14 | Day 16 | Day 18 | Day 20 | Day 22 | Day 24 |
|---|---|---|---|---|---|---|---|---|---|---|---|---|---|
| 2 | CM+AMX+PPNG | 1 | 1 | No | OK | 3.98 | 3.68 | 3.92 | 3.36 | 2.78 | 3.64 | 0.95 | 0.95 |
| 2 | CM+AMX+PPNG | 1 | 2 | No | OK | 4.38 | 0.99 | 2.99 | 0.95 | 0.95 | 0.95 | 0.95 | 0.95 |
| 2 | CM+AMX+PPNG | 1 | 3 | No | OK | 1.69 | 3.88 | 5.05 | 3.58 | 2.84 | 0.95 | 0.95 | 0.95 |
| 2 | CM+AMX+PPNG | 1 | 4 | No | OK | 5.02 | 3.40 | 1.89 | 0.95 | 0.95 | 0.95 | 0.95 | 0.95 |
| 2 | CM+AMX+PPNG | 1 | 5 | No | OK | 2.80 | 3.62 | 4.77 | 2.90 | 0.95 | 0.95 | 0.95 | 0.95 |
| 2 | CM+AMX+PPNG | 2 | 6 | No | NNG | 0.95 | 0.95 | 0.95 | 0.95 | 0.95 | 0.95 | 0.95 | 0.95 |
| 2 | CM+AMX+PPNG | 2 | 7 | No | NNG | 0.95 | 0.95 | 0.95 | 0.95 | 0.95 | 0.95 | 0.95 | 0.95 |
| 2 | CM+AMX+PPNG | 2 | 8 | No | NNG | 0.95 | 0.95 | 0.95 | 0.95 | 0.95 | 0.95 | 0.95 | 0.95 |
| 2 | CM+AMX+PPNG | 2 | 9 | No | OK | 3.25 | 3.38 | 5.33 | 4.62 | 4.26 | 1.59 | 0.95 | 0.95 |
| 2 | CM+AMX+PPNG | 2 | 10 | No | OK | 4.62 | 3.58 | 3.33 | 2.61 | 0.95 | 0.95 | 0.95 | 0.95 |
| 2 | CM+AMX+PPNG | 3 | 11 | No | OK | 5.49 | 3.60 | 4.89 | 0.95 | 0.95 | 3.64 | 0.95 | 0.95 |
| 2 | CM+AMX+PPNG | 3 | 12 | No | NNG | 0.95 | 0.95 | 0.95 | 0.95 | 0.95 | 0.95 | 0.95 | 0.95 |
| 2 | CM+AMX+PPNG | 3 | 13 | No | OK | 5.25 | 2.65 | 5.16 | 0.95 | 1.84 | 0.95 | 0.95 | 0.95 |
| 2 | CM+AMX+PPNG | 3 | 14 | No | NNG | 0.95 | 0.95 | 0.95 | 0.95 | 0.95 | 0.95 | 0.95 | 0.95 |
| 2 | CM+AMX+PPNG | 3 | 15 | No | NNG | 0.95 | 0.95 | 0.95 | 0.95 | 0.95 | 0.95 | 0.95 | 0.95 |

**Key:** CM = *Chlamydia muridarum*, AMX = Amoxicillin, NG = *Neisseria gonorrhoeae* (strain FA1090, does not produce Penicillinase), PPNG = Penicillinase-producing *Neisseria gonorrhoeae*. Experimental Groups: CM (CM-infected, not AMX-treated, not NG- or PPNG-infected); CM+AMX (CM-infected and AMX-treated, not NG- or PPNG-infected); CM+AMX+NG (CM-infected and AMX-treated, NG co-infected); CM+AMX+PPNG (CM-infected and AMX-treated, PPNG co-infected). Early Sac = sacrificed on Day 7 (Yes or No). Status: ND = not in Diestrus/Anestrus on Day 7; OK = in Diestrus/Anestrus and successfully CM and/or NG infected (included in vaginal bacterial shedding analyses); NCM = in Diestrus/Anestrus on Day 7, but CM infection not detected (excluded from vaginal bacterial shedding analyses); NNG = in Diestrus/Anestrus on Day 7, but NG or PPNG infection not detected (NNG status CM+AMX+PPNG mice were excluded from vaginal bacterial recovery analysis). Day 10-24 NG and PPNG shedding values are shown as the $\log_{10}$ of the calculated CFU/swab; limit of detection = 10, so undetectable shedding was designated as 9 CFU/swab (i.e. <10) with corresponding $\log_{10}$ value of 0.95 (indicated by red text). Shaded values are those excluded from vaginal viable PPNG recovery analysis shown in Figure 2 due to ND, or NNG status. Where no value is entered for a given Day, the mouse was sacrificed early, and thus no longer in the study on that Day. For groups CM and CM+AMX, a designation of “none” for any given Day indicates that 50 μL of vaginal swab sample was assayed on selective agar, and no contamination with either NG strain was detected.
