## Supplemental File 4 for "Murine Vaginal Co-infection with Penicillinase-Producing *Neisseria gonorrhoeae* Fails to Alleviate Amoxicillin-Induced Chlamydial Persistence"

### SUPPLEMENTAL FILE 4. Raw genital pathology data

| Expt | Group | Cage | Mouse | Early Sac | Status | Oviduct Dilation | Oviduct Inflammation | Uterus Dilation | Cervical Epithelium | Cervical Inflammation | Vaginal Epithelium | Vaginal Inflammation |
|---|---|---|---|---|---|---|---|---|---|---|---|---|
| 2 | CM | 1 | 1 | Yes | ND | 0/2 | D and O, MX++ | No | CtS | N, E++ | MISSING | MISSING |
| 2 | CM | 1 | 2 | No | OK | 0/5 | No | No | C | No | K | N, E+ |
| 2 | CM | 1 | 3 | No | OK | 0/5 | No | No | S | No | K | N, E+ |
| 2 | CM | 1 | 4 | No | OK | 0/5 | No | Yes, + | Ct | No | K | No |
| 2 | CM | 1 | 5 | No | OK | 0/5 | D, L+, F | Yes, + | MISSING | MISSING | MISSING | MISSING |
| 2 | CM | 2 | 6 | No | OK | 0/3 | D, L+, F | Yes, + | CtS | N, U+++, E+++ | K | No |
| 2 | CM | 2 | 7 | No | OK | 0/7 | D, L+, F | Yes, + | C | No | K | No |
| 2 | CM | 2 | 8 | No | OK | 0/7 | No | No | S | N, E+ | K | No |
| 2 | CM | 2 | 9 | No | OK | 0/8 | No | No | C | No | K | No |
| 2 | CM | 2 | 10 | No | OK | 0/5 | No | Yes, + | CtS | No | K | No |

| Expt | Group | Cage | Mouse | Early Sac | Status | Oviduct Dilation | Oviduct Inflammation | Uterus Dilation | Cervical Epithelium | Cervical Inflammation | Vaginal Epithelium | Vaginal Inflammation |
|---|---|---|---|---|---|---|---|---|---|---|---|---|
| 1 | CM+AMX | 1 | 1 | Yes | ND | 0/2 | D, L and M+ | No | CtS | No | K | N, E+ |
| 1 | CM+AMX | 1 | 2 | No | OK | 0/7 | No | No | PART | N, E++ | K | No |
| 1 | CM+AMX | 1 | 3 | No | OK | 0/6 | No | No | CtS | No | K | N, U++, E+++ |
| 1 | CM+AMX | 1 | 4 | Yes | ND | 1/5 | No | Yes, +++ | CtS | No | K | No |
| 1 | CM+AMX | 1 | 5 | No | OK | 0/10 | No | No | CtS | No | K | N, E++ |
| 1 | CM+AMX | 2 | 6 | Yes | ND | 0/8 | No | No | CtS | No | K | N, E +++ |
| 1 | CM+AMX | 2 | 7 | No | OK | 0/7 | No | No | CtS | No | S | N, U+ |
| 1 | CM+AMX | 2 | 8 | Yes | ND | 0/8 | No | No | C | No | K | N, E ++ |
| 1 | CM+AMX | 2 | 9 | No | OK | 0/4 | No | No | S | N, E+ | S | N, E+ |
| 1 | CM+AMX | 2 | 10 | No | NCM | 0/3 | No | No | CtS | No | S | N, E++ |
| 2 | CM+AMX | 3 | 11 | No | OK | 0/2 | No | Yes, + | C | No | K | No |
| 2 | CM+AMX | 3 | 12 | No | OK | 0/6 | D, MX++, F | No | C | No | K | No |
| 2 | CM+AMX | 3 | 13 | No | OK | 0/5 | D, L+, F | No | CtS | N, E+ | K | No |
| 2 | CM+AMX | 3 | 14 | No | OK | 0/3 | No | Yes, + | CtS | No | K | No |
| 2 | CM+AMX | 3 | 15 | Yes | ND | 0/6 | D and O, MX+++ | Yes, ++ | CtS | N, E+ | S | N, E+ |
| 2 | CM+AMX | 4 | 16 | No | OK | 0/10 | No | No | CtS | N, U+++, E+++ | K | No |
| 2 | CM+AMX | 4 | 17 | No | OK | 0/8 | No | No | C | No | K | No |
| 2 | CM+AMX | 4 | 18 | No | OK | 0/3 | No | No | CtS | N, U+++, E+++ | K | N, E+ |
| 2 | CM+AMX | 4 | 19 | Yes | ND | 0/5 | D and O, MX+++ (*) | No | CtS | No | K | No |
| 2 | CM+AMX | 4 | 20 | Yes | ND | 0/1 | D and O, MX+++ (*) | No | CtS | N, U+++, E+++ | K | N, U+++, E+ |

| Expt | Group | Cage | Mouse | Early Sac | Status | Oviduct Dilation | Oviduct Inflammation | Uterus Dilation | Cervical Epithelium | Cervical Inflammation | Vaginal Epithelium | Vaginal Inflammation |
|---|---|---|---|---|---|---|---|---|---|---|---|---|
| 2 | CM+AMX+NG | 1 | 1 | No | OK | 0/2 | D, L+, F | No | CtS | N, E+ | K | N, E++ |
| 2 | CM+AMX+NG | 1 | 2 | Yes | ND | 0/2 | D, MX+ | No | CtS | No | K | No |
| 2 | CM+AMX+NG | 1 | 3 | No | NNG | 0/6 | D, L+, F | No | CtS | N, U+++, E+++ | K | No |
| 2 | CM+AMX+NG | 1 | 4 | No | NNG | 0/4 | No | No | CtS | No | K | No |
| 2 | CM+AMX+NG | 1 | 5 | No | NNG | 0/4 | No | No | CtS | N, E+++ | K | No |
| 2 | CM+AMX+NG | 2 | 6 | No | NNG | 0/4 | No | No | CtS | No | S | No |
| 2 | CM+AMX+NG | 2 | 7 | No | NNG | 0/3 | No | No | CtS | N, E+++ | K | No |

| | | | | | | | | | | | | |
|---|---|---|---|---|---|---|---|---|---|---|---|---|
| 2 | CM+AMX+NG | 2 | 8 | Yes | ND | 0/7 | No | No | CtS | No | K | N, E+ |
| 2 | CM+AMX+NG | 2 | 9 | No | NNG | 0/1 | No | No | CtS | No | K | N, U+++, E+++ |
| 2 | CM+AMX+NG | 2 | 10 | No | OK | 0/6 | D and O, MX+ | No | CtS | N, E+++ | K | N, E+++ |
| 2 | CM+AMX+NG | 3 | 11 | No | NNG | 0/5 | No | No | CtS | No | K | N, E+++ |
| 2 | CM+AMX+NG | 3 | 12 | No | NNG | 0/4 | No | No | CtS | N, U+, E+++ | K | N, E++ |
| 2 | CM+AMX+NG | 3 | 13 | Yes | ND | 0/4 | D and O, MX+++ | No | CtS | No | K | N, U+ |
| 2 | CM+AMX+NG | 3 | 14 | No | NNG | 0/4 | No | No | CtS | No | K | N, E+++ |
| 2 | CM+AMX+NG | 3 | 15 | No | OK | 0/12 | No | No | CtS | N, E+++ | K | N, E+++ |

| Expt | Group | Cage | Mouse | Early Sac | Status | Oviduct Dilation | Oviduct Inflammation | Uterus Dilation | Cervical Epithelium | Cervical Inflammation | Vaginal Epithelium | Vaginal Inflammation |
|---|---|---|---|---|---|---|---|---|---|---|---|---|
| 2 | CM+AMX+PPNG | 1 | 1 | No | OK | 0/4 | No | No | C | No | K | No |
| 2 | CM+AMX+PPNG | 1 | 2 | No | OK | 0/3 | No | No | CtS | N, U+++, E+++ | S | N, U+, E+++ |
| 2 | CM+AMX+PPNG | 1 | 3 | No | OK | 0/2 | No | No | C | No | S | No |
| 2 | CM+AMX+PPNG | 1 | 4 | No | OK | 0/4 | No | No | CtS | No | S | No |
| 2 | CM+AMX+PPNG | 1 | 5 | No | OK | 0/2 | No | No | CtS | N, E+ | MISSING | MISSING |
| 2 | CM+AMX+PPNG | 2 | 6 | No | NNG | 3/5 | No | No | MISSING | MISSING | MISSING | MISSING |
| 2 | CM+AMX+PPNG | 2 | 7 | No | NNG | 0/4 | No | No | CtS | No | MISSING | MISSING |
| 2 | CM+AMX+PPNG | 2 | 8 | No | NNG | 0/7 | No | No | CtS | No | K | N, E+++ |
| 2 | CM+AMX+PPNG | 2 | 9 | No | OK | 0/8 | No | No | CtS | No | S | N, E+ |
| 2 | CM+AMX+PPNG | 2 | 10 | No | OK | 0/6 | No | No | CtS | No | K | No |
| 2 | CM+AMX+PPNG | 3 | 11 | No | OK | 0/8 | No | No | CtS | No | S | N, E+ |
| 2 | CM+AMX+PPNG | 3 | 12 | No | NNG | 0/9 | No | No | CtS | No | S | N, E+ |
| 2 | CM+AMX+PPNG | 3 | 13 | No | OK | 0/10 | No | No | S | N, E+ | S | N, E+ |
| 2 | CM+AMX+PPNG | 3 | 14 | No | NNG | 0/6 | D, L+, F | Yes, ++ | C | No | K | No |
| 2 | CM+AMX+PPNG | 3 | 15 | No | NNG | 0/7 | No | No | CtS | No | S | N, U+, E+++ |

**Key:** CM = *Chlamydia muridarum*, AMX = Amoxicillin, NG = *Neisseria gonorrhoeae* (strain FA1090, does not produce Penicillinase), PPNG = Penicillinase-producing *Neisseria gonorrhoeae*. Experimental Groups: CM (CM-infected, not AMX-treated, not NG- or PPNG-infected); CM+AMX (CM-infected and AMX-treated, not NG- or PPNG-infected); CM+AMX+NG (CM-infected and AMX-treated, NG co-infected); CM+AMX+PPNG (CM-infected and AMX-treated, PPNG co-infected). Early Sac = sacrificed on Day 7 (Yes or No). Status: ND = not in Diestrus/Anestrus on Day 7; OK = in Diestrus/Anestrus and successfully CM and/or NG infected; NCM = in Diestrus/Anestrus on Day 7, but CM infection not detected; NNG = in Diestrus/Anestrus on Day 7, but NG or PPNG infection not detected. No mice were excluded from pathology analyses unless tissue section was unavailable, as noted. Genital tract tissues (ovary, oviduct, uterus, cervix, vagina) were collected and stored in 4% buffered formaldehyde for 24 h followed by paraffin embedding (FFPE) according to routine procedure. From each FFPE block, 2 µm sections were cut and stained with hematoxylin and eosin. The genital sections were evaluated as complete longitudinal sections, by a board-certified pathologist. The oviduct, uterus, cervix and vagina were subjected to histologic evaluation; the ovary, present in at least one section for all mice, and similar in normal appearance across all experimental groups, was not further evaluated. For assessment of oviduct dilation, all available cross-sections were counted and the proportion of dilated/total cross-sections was recorded. Cervical epithelial appearance was recorded as columnar (C), stratified (S) or columnar to stratified (CtS). For the uterus and cervix, dilation was noted only as Yes or No, with degree of dilation recorded as mild (+), moderate (++), or severe (+++). Vaginal epithelial appearance was recorded keratinized squamous (K), or stratified (S). Presence of inflammation was recorded with no inflammation recorded as No, and with inflammatory infiltrates described qualitatively as lymphocytes (L), macrophages (M), neutrophils (N), or mixed (MX) if different inflammatory cell types were present. Inflammation localization was recorded as luminal (U) and/or intraepithelial (E) for tissues other than the oviduct, and periductal (D) and/or periovarial (O) for the oviduct. Degree of inflammation was recorded as mild (+), moderate (++), orsevere (+++); inflammation was noted as F if focal in nature for the oviduct. Missing sections (MISSING) or partially missing sections (PART) resulted in missing information, as indicated.

**Notes:**
(*) Neutrophils were also present in the oviduct lumen.
